# Cut it out: Out-of-plane stresses in cell sheet folding of *Volvox* embryos

**DOI:** 10.1101/2023.10.17.562736

**Authors:** Pierre A. Haas, Stephanie S. M. H. Höhn

**Author notes:** Joint corresponding authors.,. P.A.H. and S.M.H.H. contributed equally. The authors declare that they have no competing interests. S.M.H.H. and P.A.H. designed the study. S.M.H.H. performed experiments. P.A.H. derived the theoretical models. S.M.H.H. and P.A.H. analysed and interpreted the data and wrote the manuscript.

## Abstract

The folding of cellular monolayers pervades embryonic development and disease. It results from stresses out of the plane of the tissue, often caused by cell shape changes including cell wedging via apical constriction. These local cellular changes need not however be compatible with the global shape of the tissue. Such geometric incompatibilities lead to residual stresses that have out-of-plane components in curved tissues, but the mechanics and function of these out-of-plane stresses are poorly understood, perhaps because their quantification has proved challenging. Here, we overcome this difficulty by combining laser ablation experiments and a mechanical model to reveal that such out-of-plane residual stresses exist and also persist during the inversion of the spherical embryos of the green alga *Volvox*. We show how to quantify the mechanical properties of the curved tissue from its unfurling on ablation, and reproduce the tissue shape sequence at different developmental timepoints quantitatively by our mechanical model. Strikingly, this reveals not only clear mechanical signatures of out-of-plane stresses associated with cell shape changes away from those regions where cell wedging bends the tissue, but also indicates an adaptive response of the tissue to these stresses. Our results thus suggest that cell sheet folding is guided mechanically not only by cell wedging, but also by out-of-plane stresses from these additional cell shape changes.

---

**T**he folding of tissues into three-dimensional shapes is a crucial part of morphogenetic processes such as neurulation and gastrulation in natural embryonic development (1–4), associated birth defects such as spina bifida (5), and synthetic morphogenesis in organoids (6–9). These dramatic shape changes are regulated by a complicated interplay of mechanical forces and molecular signalling (10–17). The latter can be visualised by fluorescent tagging (18–20), but the mechanical forces must be inferred indirectly and have therefore remained one of the biggest mysteries in the life sciences.

These mechanical forces are often caused by cell shape changes, including local cell wedging through apical or basal constriction (3, 10, 21), which locally imparts a preferred, intrinsic curvature to the tissue. This intrinsic curvature drives tissue bending by generating out-of-plane forces. Such forces not only cause morphogenetic changes, but also elicit biochemical responses (22). However, physically, the mechanics and, more biologically, the role in development of different cell shape changes are still unclear: For example, what is the relative contribution to tissue folding of areas of cell wedging and adjacent areas with less pronounced cell shape changes (15)? Moreover, the local intrinsic curvature driving tissue folding is not in general compatible with the global geometry of the tissue. The folding of the tissue does not in general resolve this geometric incompatibility, so it leads to persisting stresses in the tissue termed residual stresses (23, 24). Now tissues are known to have the ability to alleviate tensile (25), compressive, or bending (26) stresses through cell movements or changes in cell adhesion or shape and this adaptation can be crucial for maintaining tissue integrity (27). How the mechanical state of the tissue is involved in this adaptation in general remains, however, an open question. Addressing these outstanding problems fundamentally relies on quantifying the spatio-temporal distribution of mechanical forces in tissues.

Over the past decade, different techniques to infer mechanical forces in tissues have therefore emerged, ranging from atomic force microscopy, traction force microscopy, and fluorescence resonance energy transfer-based molecular force sensors to optical tweezers, micropipette aspiration, magnetic beads, and liquid droplets (28, 29). However, methods for quantifying out-of-plane stresses in folding epithelia are still lacking. In fact, we are only aware of a single quantitative study of out-of-plane stresses in cell sheets: Fouchard *et al*. (26) measured the force needed to unfurl curled synthetic cell sheets and went on to estimate the active torques and the bending modulus of epithelial monolayers (30). In particular, while numerous studies have used laser ablation to infer in-plane forces from quantifications of the ensuing recoil of the tissue [see, e.g., Refs. (31–34)], extending these methods to out-of-plane forces in geometrically more complex curved dynamic tissues has proved challenging because of the difficulty of imaging out-of-plane recoils on ablation and the need for a mechanical model of these complex deformations to link the measured recoil to the mechanical forces causing it.

**Author Summary**

During the development of multicellular organisms, cell shape changes drive the emergence of complex shapes by generating out-of-plane stresses that cause tissues to fold, but quantifying such out-of-plane stresses in curved tissues has remained challenging. Here, we develop a framework combining microsurgery experiments and a mechanical model to quantify these stresses and infer mechanical properties of the tissue based on the unfurling of the tissue on being cut. We find such recoils in the alga *Volvox* and use our model to predict the stresses required to reproduce the observed tissue shape sequence quantitatively, both before and after cutting. Our results emphasise the effects of even small out-of-plane tissue stresses in development.

Here, we present our framework combining orthogonal laser ablation (OLA, Fig. 1A) and a mechanical model to quantify out-of-plane stresses and infer mechanical properties of curved cell sheets, which we apply to the gastrulation-like inversion process of the microalga *Volvox*.

**Fig. 1.**
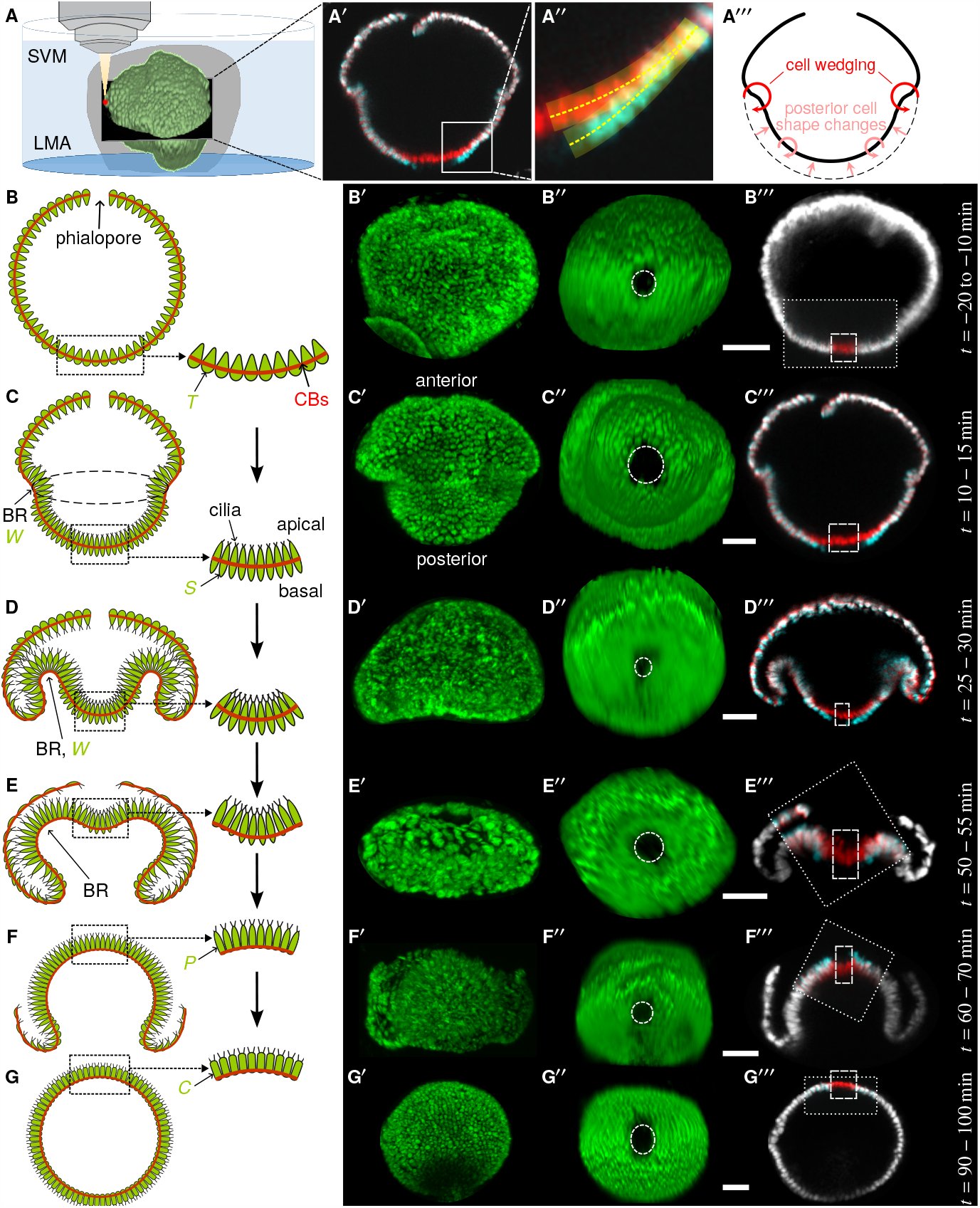
Experiments: orthogonal laser ablation (OLA) of inverting *Volvox globator* embryos. (A) Experimental setup for 2-photon microscopy and OLA, illustrated for a surface rendering of a *V. globator* embryo at an early inversion stage. Samples were embedded in low-melting point agarose (LMA), submerged in Standard *Volvox* Medium (SVM), and imaged using 2-photon microscopy. Holes were created using a separate 2-photon laser (*Materials and Methods*) at the posterior pole of embryos before, during, and after inversion. (A^*′*^) Overlay (white) of micrographs of the mid-sagittal cross-section of the embryo in panel (A), just before ablation (red) and after relaxing into its post-ablation equilibrium shape (cyan), showing the unfurled edge of the cell sheet. (A^*′′*^) Tracing of the cell sheet (transparent area) and its midline (dashed line) in the square magnified from panel (A^*′*^). (A^*′′′*^) Mechanical hypothesis: posterior cell shape changes not only contract the posterior hemisphere (45), but also induce a curvature mismatch in it that causes out-of-plane stresses distant and distinct from those generated by the wedge-shaped cells in the bend region. (B)–(G) Schematic cross-sections of *V. globator* embryos in consecutive developmental stages, modified from Ref. (41), showing associated cell shapes, the network of cytoplasmic bridges (CBs, red line), and the growing cilia. BR: bend region, W: wedge-shaped cells. Insets show cell shapes (41) near the posterior pole of the embryo (T: teardrop-shaped cells; S: spindle-shaped cells; P: pencil-shaped cells; C: columnar cells). (B^*′*^)–(G^*′*^) Maximum intensity projections of z-stacks showing lateral views of *V. globator* embryos in the developmental stages in panels (B)–(G). The anterior and posterior hemispheres are labelled in panel (C^*′*^). (B^*′′*^)–(G^*′′*^) Posterior view of the embryos in panels (B^*′*^)–(G^*′*^) after laser ablation. Dashed white line: outline of the hole created by laser ablation after relaxation into its post-ablation equilibrium shape. (B^*′′′*^)–(G^*′′′*^) Overlays (white) of micrographs of the mid-sagittal cross-section of the embryos in panels (B^*′*^)–(G^*′*^) just before ablation (red) and after relaxing into their post-ablation equilibrium shapes (cyan). Dashed lines: areas of laser ablation. Thin dotted lines in panels (B^*′′′*^), (E^*′′′*^)–(G^*′′′*^): field of view in which the laser ablation was performed. Times *t* are given relative to the start time *t* = 0 of inversion. Scale bars: 20 μm.

Inversion in *Volvox* and the related volvocine algae (Chlorophyta) is an emerging model developmental event, during which the spherical embryonic cell sheet turns itself inside out through a programme of cell shape changes (35–43). Here, we focus on type-B inversion in *V. globator* (37, 41, 44, 45, Fig. 1B–G,B^*′*^–G^*′*^). We have previously described this inversion with a mechanical model (45–47) in which the programme of cell shape changes driving the process appears as variations of the intrinsic curvatures and intrinsic stretches of an elastic shell (45, 46) and which reproduces the inversion process quantitatively (47). These *in silico* approaches have made a number of mechanical predictions (45, 47), but the elastic framework underlying them has in fact remained untested.

We start by using our ablation framework to reveal persistent residual out-of-plane stresses in the posterior hemisphere throughout inversion by showing how the cell sheet unfurls at the boundary of a circular ablation at the posterior pole. We use our mechanical model to show that the geometric incompatibilities causing this recoil result from mismatched intrinsic curvatures of the cell sheet. From quantifications of the recoil, we infer mechanical properties of the posterior hemisphere during invagination and its intrinsic curvature, which is consistent with the observed cell shapes changes. These results do not only therefore provide proof-of-principle of our ablation framework, but also test the elastic framework underlying our previous *in silico* approaches. We go on to extend our elastic model (47, 48) to reproduce the tissue shapes quantitatively at different stages of inversion. Strikingly, the resulting fitted mechanical sequence and the ablation data show a clear signature of out-of-plane stresses in the posterior hemisphere and suggest an adaptive response to these stresses. This shows that the tissue dynamics of *Volvox* inversion rely not only on cell wedging, but also on out-of-plane stresses from the additional cell shape changes in the posterior hemisphere.

**Type-B inversion in *Volvox globator*** We close this introduction with a more detailed description of type-B inversion in *Volvox globator*, summarising results that this paper relies on. After completion of an initial cell cleaveage phase (49, 50), the embryos of *V. globator* consist of a spherical monolayer of ∼3000 cells (37). Each embryo is located within a fluid-filled embryonic vesicle that is embedded in the extracellular matrix (ECM) of the parent (51–54). The embryonic cells are connected to each other by a network of cytoplasmic bridges (CBs) resulting from incomplete cytokinesis (55). A ring of cells at the anterior pole lacks CBs, leaving an opening, the phialopore. During the morphogenetic process of inversion, the embryos turn themselves inside-out through this phialopore within ∼60 −80 min in order to expose the two cilia that grow at each apical cell pole (41).

During inversion the embryonic cells, which have a microtubule-based cortical cytoskeleton, but neither cell walls nor ECM (56), remain connected to each other by CBs, so the positions of the cells relative to their neighbours are maintained (41), which suggests our elastic description of the cell sheet (45–48, 57). The global deformations of the embryonic cell sheet during inversion are driven by cell shape changes (Fig. 1B–G) that have previously been described based on electron and confocal microscopy (41).

Prior to inversion, all embryonic cells are teardrop-shaped and connected at their broadest point by CBs (Fig. 1B,B^*′*^). Type-B inversion begins (“early invagination”) with the appearance of a circular bend region (BR) just below the equator of the embryo, caused by cell-wedging and re-location of the CBs to the thinner basal cell poles (41). Simultaneously, the cells in the posterior hemisphere undergo actin-dependent thinning (58, 59) and become spindle-shaped (Fig. 1C,C^*′*^). Our previous *in silico* analyses (45, 47) indicate that active contraction of the posterior cell sheet, associated with this latter cell shape change, is necessary to explain the resulting “mushroom shape” of the embryos. As inversion proceeds (“late invagination”), a wave of cell wedging travels from the equator towards the posterior pole and the posterior hemisphere moves into the anterior one (41). The cells at the posterior pole retain their spindle shape (Fig. 1D,D^*′*^). The posterior hemisphere then moves entirely into the anterior, but a “dimple” remains where the cell sheet around the posterior pole has not yet inverted. The cells at the posterior pole remain spindle-shaped while the basal cell poles in the BR are less wedge-shaped than before, resulting in a relaxation of the curvature in the BR and near the posterior pole (Fig. 1E,E^*′*^). By the time the posterior hemisphere has fully inverted, the phialopore has widened and the anterior hemisphere has started to peel over the inverted posterior. The cells in the inverted posterior have adopted a pencil shape (Fig. 1F,F^*′*^) at this stage. Closure of the phialopore marks the completion of inversion and the end of embryogenesis in *Volvox*. At this stage, all cells are pencil-shaped with pointy apical cell poles (41). Within 30 − 60 min after inversion, all cells shorten, widen, and adopt a columnar shape while the radius of the juvenile spheroid increases (Fig. 1G,G^*′*^).

## Experiments

### Orthogonal laser ablation (OLA) in type-B *Volvox* inversion

To quantify out-of-plane stresses in *Volvox* inversion, we performed laser ablation experiments on *V. globator* embryos in consecutive stages of inversion (*Materials and Methods*). In order to capture out-of-plane elastic responses to these ablations, they were performed on the mid-sagittal cross-sections of the axisymmetric *V. globator* embryos. Cross-sections were imaged using 2-photon microscopy (Fig. 1A and *Materials and Methods*), and approximately circular holes were created at the posterior pole of the embryos by laser ablation using a separate 2-photon laser (*Materials and Methods*). The axisymmetry of the embryos and the ablations ensures that any subsequent deformation of the cell sheet is approximately axisymmetric, too. The dimensions of the resulting hole in the cell sheet were determined in three-dimensional datasets recorded subsequently (*Materials and Methods*).

### Observations

Laser ablations were performed at the posterior pole of embryos in consecutive developmental stages before, during, and after inversion (Fig. 1B^*′′*^–G^*′′*^,B^*′′′*^–G^*′′′*^).

Prior to inversion, embryos did not show any recoil on laser ablation (Fig. 1B^*′*^–B^*′′′*^), i.e. no deformation within at least 2 min following ablation (*N* = 5 embryos), indicating that the cell sheet is not residually stressed before the onset of inversion. However, during early invagination, ablation led to outwards unfurling of the edges of the cell sheet and relaxation into a new equilibrium shape (Fig. 1C^*′′*^,C^*′′′*^; *N* = 14), showing that out-of-plane residual stresses have appeared in the cell sheet. Ablations during the late invagination stage again led to unfurling of the edges of the cell sheet (Fig. 1D^*′′*^,D^*′′′*^; *N* = 5). When ablations were performed at the “dimple stage” (Fig. 1E,E^*′*^), the unfurling cell sheet adopted a curvature closer to that in the BR (Fig. 1E^*′′*^,E^*′′′*^; *N* = 5), indicating a mechanical effect of the BR on the post-ablation equilibrium shape. Interestingly, for ablations at the stage where the posterior hemisphere has fully inverted (Fig. 1F,F^*′*^), the direction of unfurling changed compared to earlier timepoints (Fig. 1F^*′′*^,F^*′′′*^; *N* = 7). Finally, for posterior ablations after the cells have become columnar (Fig. 1G,G^*′*^), the cell sheet no longer shows any recoil within at least 2 min after laser ablation (Fig. 1G^*′′*^,G^*′′′*^; *N* = 4), indicating that the residual stresses have relaxed.

At all stages of inversion observed, unfurling of the edges of the cut upon laser ablation and relaxation into the postablation equilibrium shape took approximately 4 −10 s. Within this time, the embryos otherwise maintained their shape: Morphogenetic inversion movements, i.e. global embryonic shape changes distinct from the local ablation response, were only distinguishable after at least 1 min. While the hole caused by laser ablation did not close up, the embryos did complete inversion after ablation, unless the cut size exceeded about half the radius of the posterior hemisphere in which case the anterior hemisphere failed to invert (not shown).

The observed outward unfurling of the cell sheet suggests that the preferred, intrinsic curvature of the cell sheet differs from the curvature into which its unablated spherical shape forces the cell sheet. We therefore hypothesise that such a curvature mismatch gives rise to out-of-plane stresses in the posterior hemisphere, including stresses distant and hence distinct from the out-of-plane stresses in the bend region of wedge-shaped cells (Fig. 1A^*′′′*^). The thinning of cells from teardrop to spindle shapes driving contraction of the posterior hemisphere also changes its curvature (46, Fig. 1A^*′′′*^,B,C), so would contribute to this mismatch of curvatures. The change of the direction of unfurling with the change of the sign of the curvature of the posterior hemisphere (Fig. 1B^*′′′*^,F^*′′′*^) is consistent with this hypothesis if the intrinsic curvature of the tissue is smaller in magnitude than its actual curvature.

### Origin of out-of-plane stresses in *Volvox* inversion

#### Mechanical toy problems

To test this hypothesis, we need to understand the mechanics of ablations in curved tissues. For this purpose, we introduce three mechanical toy problems.

#### Toy problem 1: deformations of a spherical elastic shell with mismatched intrinsic curvatures

Our first toy problem studies the effect of curvature mismatch on cell sheet shapes before ablation: we consider a complete spherical shell of unit radius and (relative) thickness *h* ≪1. The intrinsic curvatures of the shell are *κ*^0^ = *k* ≠ 1, different from the undeformed curvature of the shell. This curvature mismatch causes the shell to deform; we assume that the shell remains spherical, and denote by *f* its deformed radius (Fig. 2A).

**Fig. 2.**
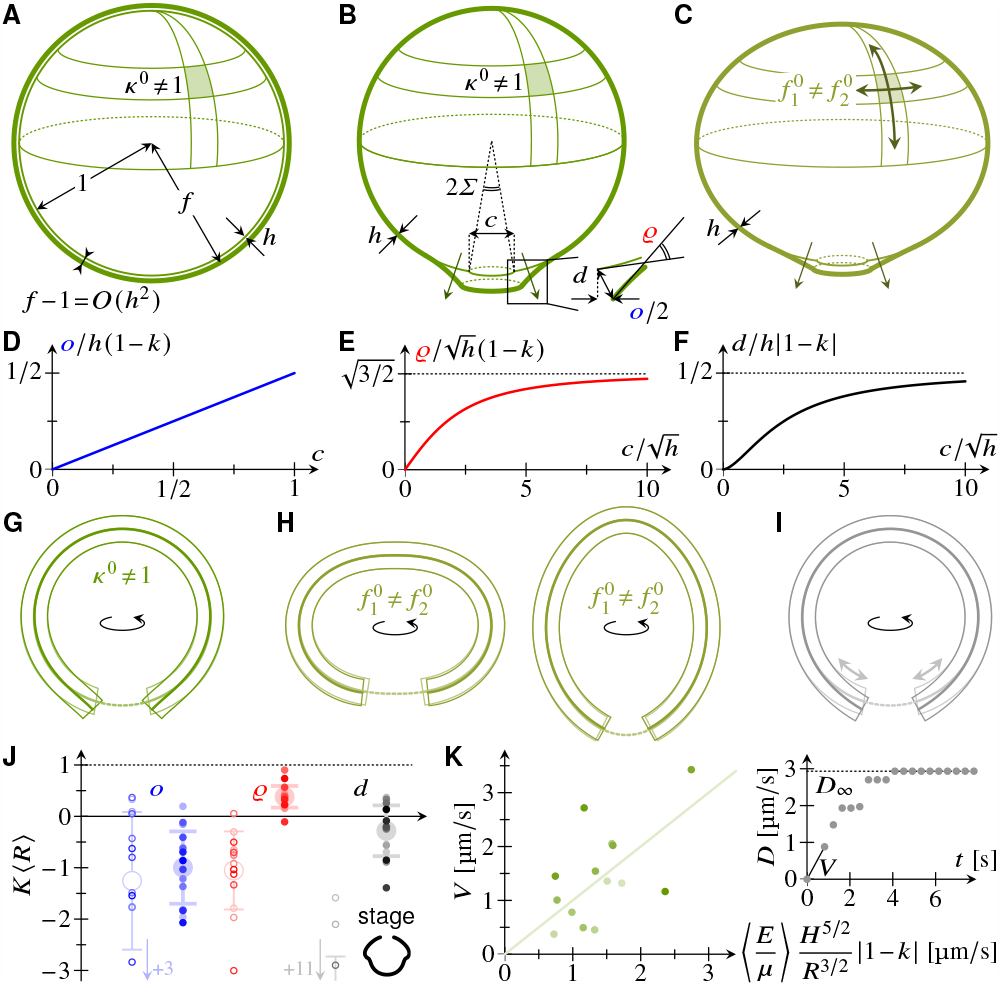
Origin of out-of-plane stresses in *Volvox inversion*: mechanical toy problems and analysis of experiments. (A) Toy problem 1: a spherical shell of unit radius and thickness *h* ≪ 1 deforms to a radius *f* = 1 + *O*(*h*^2^) due to mismatched intrinsic curvatures *κ*^0^ = *k*⁄= 1. (B) Toy problem 2: this mismatch of intrinsic curvatures causes a circular cut of angle 2*Σ* and radius *c* = 2 sin *Σ* to open. Inset: definition of the opening *o*, rotation *ϱ*, and displacement *d* of the cut edge. (C) Toy problem 3: anisotropic contraction also causes recoil on ablation. (D) Plot of *o* ∼ *h*(1 −*k*) against *c*. (E) Plot of asymptotic approximation for 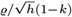 against 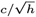 lot of asymptotic approximation for *d/h*(1 − *k*) against 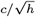. (G) Numerical solution of toy problem 2: plot of cross-sections preablation (light) and post-ablation (dark), showing a recoil qualitatively consistent with experiments. (H) Numerical solutions of toy problem 3: the recoil is much smaller than in experiments despite the excessive anisotropy. (I) Fast localised relaxation can also reproduce the experimental recoil qualitatively. (J) Estimated dimensional intrinsic curvatures *K*, nondimensionalised with the average posterior radius ⟨*R*⟩ during invagination. Estimates computed from asymptotic results (open markers) and numerical solutions of toy problem 2 (filled markers) using measurements of *o, ϱ, d* for *N* = 14 *Volvox* embryos at early invagination (inset). Arrows: off-scale values; large markers and error bars: mean and standard deviation of estimates. (K) Plot of the corresponding initial recoil velocity *V* against its scaling. Variables: elastic modulus *E*, effective viscosity *μ*, (dimensional) cell sheet thickness *H*, posterior radius *R* during invagination, (dimensionless) estimated intrinsic curvature *k*. A straight-line fit estimates the mean ⟨*E/μ*⟩. Inset: example measurement of the (dimensional) cut displacement *D* against time, showing the initial velocity and equilibrium displacement *D*_*∞*_ used for estimates.

This radius is determined by the competition of the strains *E* = *f* − 1 due to the stretching of the shell and the bending strains *K* = 1*/f* −*k* resulting from the difference of the actual and intrinsic curvatures of the shell. The elastic energy of the shell is the sum of its stretching and bending energies (60), so is proportional to

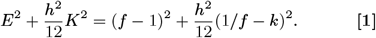

Assuming *k* = *O*(1), this is minimised for

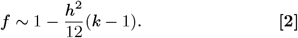

As expected, the shell grows (*f >* 1) or shrinks (*f <* 1) if its preferred curvature is flatter (*k <* 1) or larger (*k >* 1) than its undeformed curvature, but the deformations are asymptotically small compared to the shell thickness.

There is an interesting mechanical subtlety of this calculation: Though this argument gives the correct *O*(*h*^2^) scaling for the deformations of the shell, the smallness of the deformations is beyond the realm of validity of the shell theory assumed in writing down Eq. **1**. For this reason, the correct prefactor has to be calculated within bulk nonlinear solid mechanics; the calculation is given in *SI Appendix*, where we also discuss some of the suprising behaviour revealed by that prefactor which shows how even seemingly innocuous problems in nonlinear mechanics, such as this one, can break our intuition.

#### Toy problem 2: circular ablation of a spherical elastic shell with mismatched intrinsic curvatures

Next, we study the mechanics of a circular ablation in a shell with mismatched intrinsic curvatures. The mismatched intrinsic curvatures and the no-torque boundary condition at the rim of the cut cause the shell to deform near the cut. For an ablation of angle 2*Σ*, i.e. radius *c* = 2 sin *Σ* (Fig. 2B), asymptotic solution of the equations of shell theory for *h* ≪1 (*SI Appendix*) shows that the cut opens by an amount (Fig. 2D)

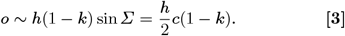

Importantly, this is asymptotically larger than the deformation without ablation described by Eq. **2**. Asymptotic approximations can also be found (*SI Appendix*) for the rotation *ϱ* of the rim of the cut and its displacement *d* due to the ablation (Fig. 2E,F).

#### Toy problem 3: circular ablation of a spherical elastic shell with anisotropic contraction

Opening of an ablation does not require intrinsic curvature mismatches, however: As explained in more detail in *SI Appendix*, geometry requires the stretches of an axisymmetric spherical shell to be equal at its poles, so anisotropic contraction will cause stresses there and hence result in a recoil on ablation (Fig. 2C).

### Discussion

These results allow us to test our hypothesis, that there is a curvature mismatch during *Volvox* inversion. First, toy problems 1 and 2 emphasise that ablation experiments are needed: The unablated deformations are too small to be visualised in experiments; only the deformations from ablations are comparable to the thickness of the cell sheet, so allow quantification of curvature mismatches that cannot be quantified from the unablated cell sheet only.

### The observed recoil during Volvox invagination results from a curvature mismatch

Next, solving for post-ablation shapes in toy problems 2 and 3 numerically (*Materials and Methods*), we observe that the recoil for an intrinsically flat shell (*k* = 0) agrees with experiments qualitatively (Fig. 1B^*′′′*^, Fig. 2G). By contrast, the recoil resulting from anisotropic contraction is much smaller than the experimentally observed one, even for blatantly excessive anisotropies (Fig. 1B^*′′′*^, Fig. 2H). This does not yet allow the conclusion that the observed recoil results from a curvature mismatch: Indeed, fast biological processes could *a priori* be triggered close to the ablation site to lead to a fast active relaxation of the shell there. Indeed, this could result in a geometric incompatibility leading to a recoil qualitatively consistent with experiments (Fig. 2I and *Materials and Methods*). If however this mechanism underlied the experimental observations, we would expect to observe a recoil for preand post-inversion ablations, too, which we do not (Fig. 1B^*′′′*^,G^*′′′*^). We therefore discard this possibility, and conclude that the observed recoil is only consistent with a curvature mismatch that existed before ablation.

### Quantitative estimates of the curvature mismatch during Volvox invagination are consistent with the observed cell shape changes

To make this analysis more quantitative, we measured the recoil on ablation in *N* = 14 *Volvox* embryos during early invagination (Fig. 1C–C^*′′′*^ and *Materials and Methods*), and extracted estimates of intrinsic curvatures (Fig. 2J) from the measurements of the cut opening *o*, rotation *ϱ*, and displacement *d* using the asymptotic approximations (Fig. 2D–F) and numerical solutions of toy problem 2. The hemispherical shape of the posterior at these early inversion stages (Fig. 2J, inset) justifies using toy problem 2 for this inference. The estimates based on asymptotic approximations have a much larger variance than those based on numerical solutions (Fig. 2J). This suggests that, while the asymptotic estimates are useful to understand the mechanical basis of the recoil upon ablation, the cell sheet of *Volvox* is too thick for them to yield good quantitative estimates. Meanwhile, we observe that the estimate, based on numerics, of mean intrinsic curvature from *o* has a larger standard deviation than the estimates from *ϱ* and *d* (Fig. 2J). Moreover, the latter are within one standard deviation of each other, while the former is not (Fig. 2J). This is perhaps not unexpected, as measuring *o* requires quantifying smaller deformations. We therefore estimate the mean intrinsic curvature of the cell sheet based on *ϱ* and *d* only, finding ⟨*k*⟩ ≈ 0.05 ≪ 1. Importantly, this estimate is consistent with the observed cell shape changes: The spindle-shaped cells in the posterior during early *Volvox* invagination are symmetric, and connected at their midplane by cytoplasmic bridges (Fig. 1C, inset), suggesting *k* = 0. By contrast, the teardrop-shaped cells in preinversion embryos (Fig. 1B, inset) suggest a positive intrinsic curvature, and indeed the absence of recoil observed preinversion (i.e. the absence of residual stresses) requires *k* = 1.

### Recoil velocity measurements yield a bound on the elastic modulus of the cell sheet

The (dimensional) recoil velocity *V* is set by the interplay between the elastic force driving it and viscous dissipation in the tissue and surrounding fluid. A scaling argument (*SI Appendix*) yields

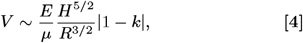

in which *E* is the elastic modulus of the cell sheet, *μ* is an effective viscosity, and *H* and *R* are the dimensional thickness and posterior radius of the cell sheet. We obtain *V* from measurements of the initial recoil velocity of the cell sheet (Fig. 2K). We stress our use of this initial velocity for these dynamic measurements, by contrast with the above matching of (quasi-)equilibrium quantifications of the recoil to a static toy problem (Fig. 2K, inset). From the scaling relation, we obtain a mean value ⟨*E/μ*⟩ ≈ 0.6 s^*−*1^ (Fig. 2K). With the lower bound *μ* ≳ 0.8 mPa s corresponding to the viscosity of algal growth medium (61), so neglecting dissipation within the tissue, this yields *E* ≳ 0.5 mPa, much lower than the values reported for confluent tissues, yet perhaps appropriate for an extremely floppy, non-confluent tissue. With *H* ≈ 11 μm (41), the bending modulus *K* = *EH*^3^ ≳ 7 10^*−*19^ J, well above the energy of thermal fluctuations, provides a sanity check of this estimate.

### Effect of out-of-plane stresses in *Volvox* inversion

Having thus shown that out-of-plane stresses resulting from a curvature mismatch persist during *Volvox* invagination, we ask: What is the effect of these out-of-plane stresses on the tissue shape sequence during inversion?

#### Quantitative model of *Volvox* inversion

The toy models in the previous section can describe the shapes of *Volvox* embryos during early invagination locally, close to the posterior ablation, but cannot capture the global embryo shapes and mechanics. We therefore extended our previous detailed morphoelastic model of the cell shape changes of inversion (47) and fitted parameters encoding the cell shape changes driving inversion (41) to the average cell sheet midlines at various timepoints of inversion (47), as described in detail in *Materials and Methods* and *SI Appendix*. We then performed numerical ablations on these fitted shapes and compared *in vivo* and *in silico* results (Fig. 3A–M).

**Fig. 3.**
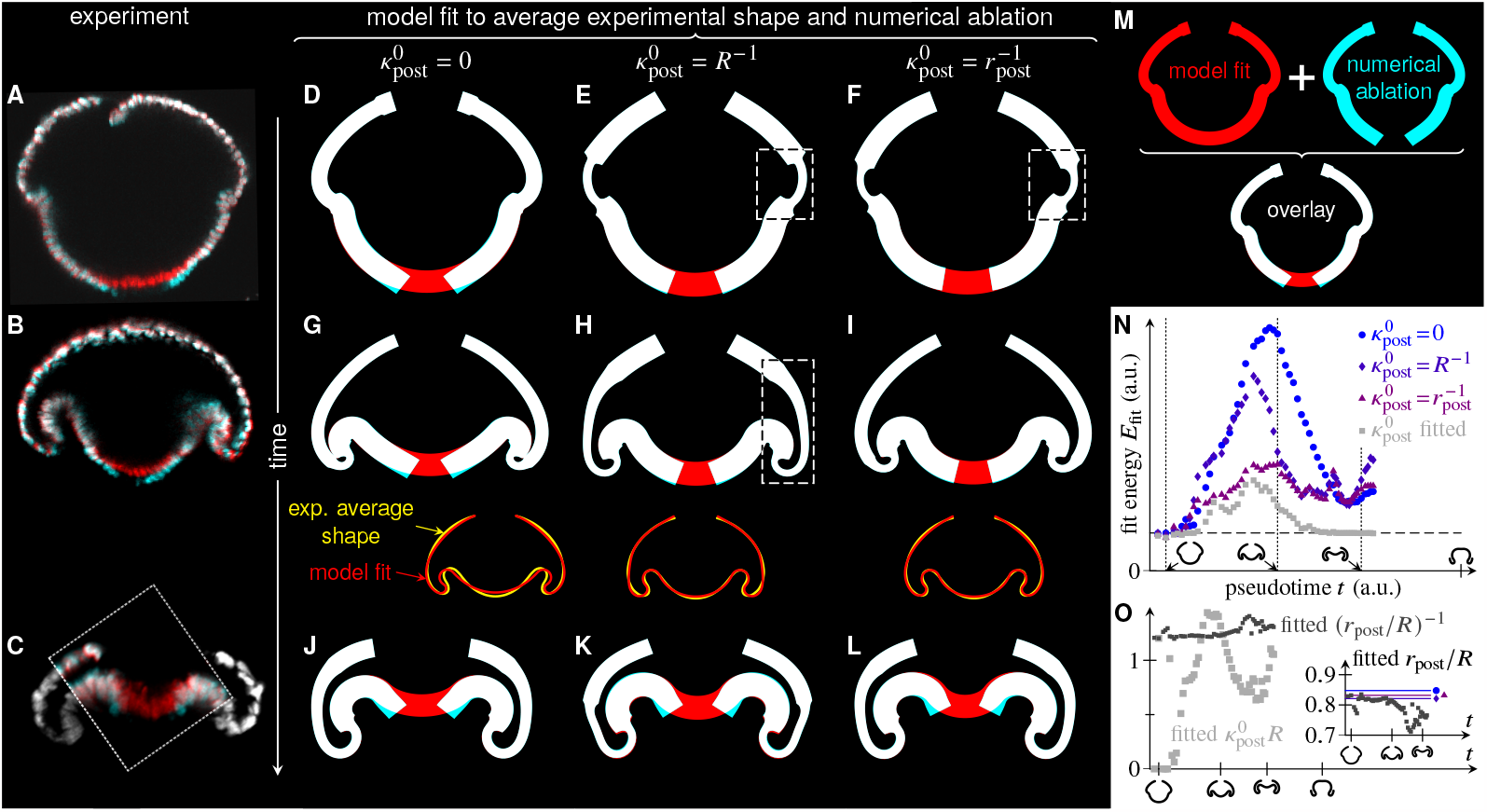
Effect of out-of-plane stresses in *Volvox* inversion: fits of the elastic model to average experimental cross-sections (47) and numerical ablations. (A) Overlay of experimental cross-sections (white) before (red) and after (cyan) a posterior ablation at an early invagination stage (Fig. 1B^*′′′*^). (B) Analogous plot for an ablation at a late invagination stage (Fig. 1C^*′′′*^). (C) Analogous plot for an ablation at a mid-inversion stage (Fig. 1D^*′′′*^). Outside the dotted rectangle, only pre-ablation shapes are plotted (white). (D) Fit, for zero posterior intrinsic curvatures 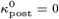, of the elastic model to the average experimental cross-section at an inversion timepoint corresponding to panel (A) (red), overlaid with equilibrium shape after numerical ablation (cyan), showing a recoil on ablation. (E) Analogous plot, for posterior intrinsic curvatures 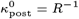, equal to the undeformed curvature of the embryo. The recoil is reduced. The dashed box highlights thinning of the cell sheet in the fitted shape. (F) Analogous plot, for posterior intrinsic curvatures 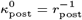, equal to the inverse contracted intrinsic radius of the posterior. There is almost no recoil. (G) Fit, for 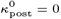, of the elastic model to the average experimental cross-section at an inversion timepoint corresponding to panel (B), and numerical ablation. Inset: comparison of fitted shape and average experimental cross-section. (H) Analogous plot for 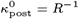. The fitted shape reproduces the experimental shape better than that in panel (G). (I) Analogous plot for 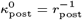. The fit is again better than that in panel (G). (J) Fit, for 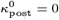, of the elastic model to the average experimental cross-section at an inversion timepoint corresponding to panel (C), and numerical ablation. (K) Analogous plot with 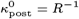. The recoil on ablation is qualitatively similar. (L) Analogous plot with 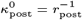. (M) Illustration of the numerical ablation plots using the plots in panel (D): the superposition of the fitted shape before numerical ablation (red) and that after numerical ablation (cyan) defines the overlaid shape (white). (N) Plot of the fit energy *E*_fit_ (arbitrary units) against inversion (pseudo)time for fits with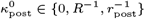 and for a fit in which 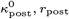 are among additional fitting parameters. *E*_fit_ = 0 would indicate a perfect fit; the horizontal dashed line indicates the value of *E* at which fitting was cut off. Pseudotimepoints used for the fits in panels (D)–(L) are highlighted. (O) Plots of fitted 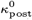 and (*r*_post_*/R*)^*−*1^ against inversion (pseudo)time. Inset: plot of fitted *r*_post_*/R* against inversion pseudotime. The constant values used for the fits with 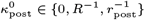 are highlighted.

#### Results for early invagination stages are consistent with the observed cell shape changes and the estimates from ablation recoils

We first considered an early invagination timepoint (Fig. 3A), for which we fitted the average embryo shape for three different values of the posterior intrinsic curvature 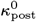 (Fig. 3D–F): 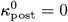, corresponding to the flat posterior intrinsic curvature predicted by the quantitative estimates of the previous section and the observed cell shape changes in the posterior; 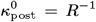, equal to the inverse radius of the undeformed embryo, i.e. the value suggested at pre-inversion stages by the lack of recoil on ablation; 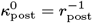, equal to the inverse intrinsic contracted radius of the posterior resulting from the cell shape changes to spindle-shaped cells (Fig. 1C). With this last value, the contracted shape of the uninverted posterior is unstressed. The model can reproduce the average embryo midlines well for all three values of 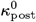 (Fig. 3D–F), although the fit parameters inferred from the cell sheet midlines for the latter two values imply a thinning of the anterior hemisphere of the cell sheet (highlighted in Fig. 3E,F) that is only observed experimentally at slightly later stages of invagination, when it associated with the formation of pancake-shape cells there (41, Fig. 1D). However, numerical ablations reveal that these three fitted shapes are mechanically different (Fig. 3D–F): Only the first case leads to a sizeable recoil on ablation, qualitatively consistent with experiments. The fitting thus suggests that 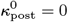 and is therefore consistent with the estimates of the previous section and the observed cell shape changes.

#### Results for late inversion stages are consistent with the observed expansion of the region of cell wedging

Next, we considered a late inversion timepoint (Fig. 3C), around the time of phialopore opening. Again, all three values of posterior intrinsic curvature could reproduce the observed cell sheet midlines (Fig. 3J–L) and the fitted shapes even reproduce the observed thinning of the anterior hemisphere (41) even though the fits are based on the cell sheet midlines only. Moreover, numerical ablations lead to similar recoils in all three cases (Fig. 3J–L). This is consistent with the observed expansion of the wave of cell wedging towards the posterior pole (41): The recoil magnitude is set no longer by the geometric incompatibilities of the few remaining spindle- or pencil-shaped cells (Fig. 1E, inset), but by that of the wedge-shaped cells in the bend region (Fig. 1C).

## Discussion

These results thus provide a check of the fitting and suggest that the mechanics of out-of-plane stresses in *Volvox* inversion can be captured correctly by this fitting. In turn, this indicates that we can use the fit results to understand out-of-plane stresses at other stages of inversion.

### The fitted mechanics at midinversion stages predict changing posterior out-of-plane stresses

We thus turn to a midinversion stage next (Fig. 3B), again fitting the model to the experimental data for the three different values of 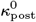 (Fig. 3G–I). Interestingly, the quality of the fit to the average experimental inversion shape is considerably worse for 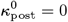 (Fig. 3G, inset) than for the other two values (Fig. 3H–I, insets). Interestingly, the fit parameters inferred from the cell sheet midlines in the case 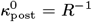 imply a thickness gradient in the anterior hemisphere (Fig. 3H) that is consistent with the formation of pancake-shaped cells starting in the anterior fold and propagating towards the phialopore, as described previously (41, Fig. 1D). All of this suggests that the intrinsic curvature of the spindle-shaped cells and hence the out-of-plane stresses in the uninverted posterior change at midinversion stages. The fact that a better fit (Fig. 3H–I) is obtained for positive values of that are associated with reduced geometric incompatibilities in the uninverted posterior begs the question: Could this change constitute an adaptive response of the tissue to the stresses resulting from the geometric incompatibility generated by the spindle-shaped cells in the posterior?

### The fitted mechanical sequence hints at an adaptive response to out-of-plane stresses

To provide a hint of an answer to this question, we considered fits of our model to the experimental data for more developmental timepoints before phialopore opening^∗^ and for the three values of 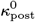. While the fit energies (Fig. 3N) are comparable at early and late stages of inversion, those at mid-inversion stages are lowest, by some margin, for the fit with 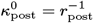. We have already noted above that the contracted shape of the uninverted posterior is unstressed in this case. This therefore hints that 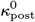 changes in such a way as to relieve the geometric incompatibilities in the uninverted posterior, and hence at an adaptive response to the stresses. To substantiate this hint, we performed a further fit in which 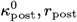 are among additional fitting parameters (*SI Appendix*). The resulting fit energy (Fig. 3N) is not substantially lower than that of the more constrained fit with 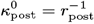 at mid-inversion stages. Moreover, the fitted values of 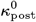 (Fig. 3O) increase from zero at the start of invagination to values comparable to *r*^*−*1^ at mid-inversion stages. All of this lends further support to the hypothesis of an adaptive response to stresses in the tissue. Interestingly, the fitted values of 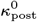 (Fig. 3O) decrease and those of *r*_post_ (Fig. 3O, inset) decrease before increasing again at mid-to-late inversion stages, when the fit energy of this fit becomes rather smaller than that of the other fits (Fig. 3N). All of this could be a signature of the pencil-shaped cells (41, Fig. 1E, inset) that begin to form at these stages of inversion (and another adaptive response): Indeed, the position of cytoplasmic bridges in these cells may suggest 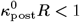, and they are narrower than spindle-shaped cells (41), indicating reduced *r*_post_. We discuss these results further in *SI Appendix*.

## Conclusion

We have combined ablation experiments and a detailed elastic model to establish the origin and analyse the mechanical effects of out-of-plane stresses during *Volvox* inversion. Strikingly, our results show how changes of posterior intrinsic curvature— albeit small compared to the intrinsic curvature imposed by the wedge-shaped cells in the bend region—contribute to the observed tissue shape sequence of *Volvox* inversion. In particular, the theory predicts an intriguing adaptive response of the geometric incompatibilities in the cell sheet to its out-of-plane stresses.

Tissue bending is not therefore all about apical constriction: Rather, out-of-plane stresses from other cell shapes changes also contribute mechanically to *Volvox* inversion. This is possible for biological tissues are not infinitely thin: If they were, the asymptotic separation of stretching and bending energies (60) would imply that only the large bending deformations of apical constriction (30, 48) that break this asymptotic separation would have an effect comparable to that of in-plane stretching deformations. The question now becomes: Do these out-of-plane stresses that impact the shapes of inverting *Volvox* embryos and their adaptive response predicted by theory have a mechanical function? Since inversion is still possible for all three scenarios in Fig. 3D–L, they are not required mechanically for inversion, unlike the formation of wedge-cells or contraction, which is known to be required mechanically for the closely related type-A inversion in *V. carteri* (57–59). However, one might speculate that these stresses or equivalently the stored elastic energy contribute to the mechanical robustness of *Volvox* inversion. Also, these stresses and their predicted adapation during inversion could have a function related to mechanosensing and hence to the unknown signalling processes that orchestrate inversion. Testing such hypotheses will require again close integration of experiment and theory and, in particular, dynamic quantification of cell shapes to link “microscopic” cell-level changes to geometric incompatibilites and hence stresses at the “macroscopic” tissue level.

Our experimental and theoretical approach sets out a framework for inferring the mechanical state and properties of curved tissues based on their unfurling on ablation, which we expect to be applicable to a broad range of problems in cell sheet folding. However, it has also revealed key differences between the analysis of ablations in curved and flat tissues: First, the standard measure of recoil on ablation in flat tissues, i.e. what we termed the cut opening above, may not be appropriate for analysing ablations in curved tissues, because it requires quantification of smaller deformations than the measures of cut rotation and cut edge displacement that we introduced here, and is therefore more prone to experimental noise. Second, the inference of cell sheet properties from the deformations on ablations is more complex in curved tissues: It may require numerical solution of a “toy problem” rather than use of a closed-form expression because tissues are not asymptotically thin, as already noted above: The tissue thickness, which is the natural small parameter for asymptotic solution of such toy problems, may not be sufficiently small for closed-form asymptotic expressions to enable quantitative inference.

Finally, from a more physical point of view, it is striking that the “simple” tissue shapes of early *Volvox* invagination did not constrain the mechanical state of the cell shape to the same extent as the more complex shapes at mid-inversion stages: At early invagination stages, knowledge of the unfurling of the tissue on ablation was required to distinguish between the three mechanical scenarios discussed in Fig. 3D–F, while, at later stages, we could distinguish between these possibilities based on the shapes of cell sheet alone (Fig. 3G–I). It is tempting to ask whether this observation generalises: Do more complex tissue shapes intrinsically contain more information about the mechanical state of the tissue? Of course, this begs a more basic question: What is the right way of quantifying tissue shape complexity? Addressing these question both in simple physical model problems and in simple biological systems like *Volvox* or synthetic, organoid models will allow us to take the first steps towards addressing, too, the much more fundamental biological problem that is the relation between tissue shape (complexity) and biological function.

## Materials and Methods

### Model organism cultivation

Wild-type strain *Volvox globator* Linné (SAG 199.80) was obtained from the Culture Collection of Algae at the University of Göttingen, Germany (62), and cultured as described previously (45, 63) in liquid Standard *Volvox* Medium (SVM) with a cycle of 16 h light at 24^*°*^C and 8 h dark at 22^*°*^C. *V. globator* cultures in the asexual life cycle of these monoecious microalgae were used.

### 2-photon microscopy and laser ablation experiments

*Volvox* spheroids containing embryos undergoing inversion were embedded in 2 μl of 1% low-melting-point agarose (LMA), covered with SVM, and imaged using a Trim Scope 2-photon microscope (LaVision). A laser line at *λ* = 1040 nm was used for imaging and a separate laser line at *λ* = 900 nm for performing laser ablations. As a single chloroplast mostly fills each *Volvox* cell, chlorophyll-autofluorescence was detected at *λ >* 647 nm, and used for visualising the embryonic cell sheet. Z-stacks were recorded before and after each laser ablation experiment, with a z-step of 2 μm. To maximise acquisition speed and capture the elastic response to the laser cut, a single plane was imaged during ablation experiments, which removed cells within a radius of 3 − 6 μm of the posterior pole of the cell sheet. Videos of the mid-sagittal plane were recorded at 1 − 4.5 fps for at least 10 s before and 30 s after laser ablation. Videos used for determining velocities of elastic recoils were recorded at the maximal speed possible for the field of view size used, corresponding to 2.5 *−* 4.5 fps.

### Analysis of experimental data

Outlines of the posterior hemisphere were traced manually on recordings of midsagittal embryo crosssections using Fiji (64). To trace the midline of the cell sheet, the line width was set to fit the thickness of the cell sheet. Recoil quantification was then performed using custom Matlab (The MathWorks, Inc.) scripts; in particular, the fit in Fig. 2K uses the polyfitZero function by M. Mikofski, obtained from the Matlab file exchange (file 35401). Posterior cell sheet radii were estimated by fitting a circle to the cross-sections in Fiji. Cell sheet thicknesses were estimated by scaling the chloroplast thicknesses measured in Fiji so that the resulting mean value matched that reported previously (41).

### Morphoelastic shell theory

To describe *Volvox* morphogenesis in terms of the axisymmetric deformations of an elastic shell with varying intrinsic stretches and curvatures, we use the shell theory derived in the biologically relevant limit of “large bending deformations” in Ref. (48): For an axisymmetric elastic shell of relative thickness *h*, we denote by *s* and *S* the respective arclengths of the undeformed and deformed cross-sections of the axisymmetric shell. These cross-sections are described by their distances *ρ, r* from the axis of symmetry and and the tangent angle *ψ* of the deformed cross-section. The meridional and circumferential stretches and curvatures of the deformed shell are thus

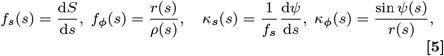

and we denote by 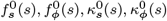 their intrinsic, preferred values. The differences between the actual stretches and curvatures and their intrinsic values define the shell strains

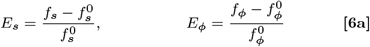

and curvature strains

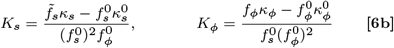

that appear in the elastic energy

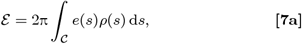

where the integration is along the cross-section 𝒞 of the shell, and the energy density is

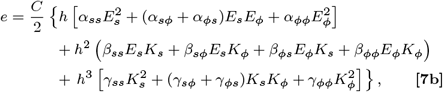

in which *C* is a material parameter and *α*_*ss*_, *α*_*sϕ*_, *α*_*ϕs*_, *α*_*ϕϕ*_, *β*_*ss*_, *β*_*sϕ*_, *β*_*ϕs*_, *β*_*ϕϕ*_, *γ*_*ss*_, *γ*_*sϕ*_, *γ*_*ϕs*_, *γ*_*ϕϕ*_ are functions of 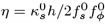 only, the (complicated) explicit expressions for which are given in Eqs. (58) and (64) of Ref. (48).

To find the deformed shape of the shell resulting from the imposed intrinsic stretches and curvatures numerically, we solve the boundary value problem associated with derived in Ref. (48) using the bvp4c solver of Matlab.

### Numerical solution of the toy problems

We solve these equations for a circular cross-section *ρ*(*s*) = sin *s* for *Σ* ⩽ *s* ⩽ π. At *s* = π, we impose the boundary conditions *r* = *ψ* = 0; if *Σ >* 0, we impose no-force and no-torque conditions at *s* = *Σ* which, from Ref. (48), take the form *α*_*ss*_*E*_*s*_ + *α*_*sϕ*_*E*_*ϕ*_ + *h*(*β*_*ss*_*K*_*s*_ + *β*_*sϕ*_*K*_*ϕ*_) = 0, *β*_*ss*_*E*_*s*_ + *β*_*ϕs*_*E*_*ϕ*_ + *h*(*γ*_*ss*_*K*_*s*_ + *γ*_*sϕ*_*K*_*ϕ*_) = 0; if *Σ* = 0, the boundary conditions at *s* = 0 are instead *r* = *ψ* = 0.

For toy problem 2, we take 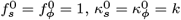. For toy problem 3, we take 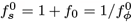, where is a constant that expresses the anisotropy of contraction, and 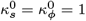. Finally, to describe a possible fast active deformation of the tissue on cutting, we 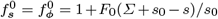 if *Σ* ⩽ *s* ⩽ *Σ*+*s*0for some length scale *s*_0_,and 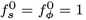 otherwise, as well as 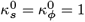 results do not seem to depend strongly on the form of the decay of away 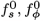 from *s* = *Σ* (not shown). The parameter values for Fig. 2G–I are *h* = 0.3, *Σ* = 0.3, *k* = 0, *f*_0_ = ± 0.1, *F*_0_ = 0.35, *s*_0_ = 0.2.

### Fitting the morphoelastic model to the experimental data

For a quantitative description of *Volvox* inversion, similarly to Ref. (47), we consider again a circular cross-section *ρ*(*s*) = sin *s*, but now for *Σ* ⩽ *s* ⩽ π − *P*, where *P* is the phialopore size. We set *P* = 0.3 and *h* = 0.15, as in Ref. (47). The boundary conditions are those of the toy problems discussed above, except for *s* = π − *P*, where we impose no-force and no-torque conditions.

We define functional forms for the intrinsic stretches and curvature functions 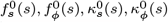 that allow a minimal representation of the cell shape changes observed during *Volvox* inversion (41), described in detail in *SI Appendix*. These functional forms define a number of fitting parameters, which, with *Σ* = 0, were fitted to the average inversion shapes (47), by minimising a fit energy *E*_fit_ that measures the difference between the average and fitted shapes using the Matlab function fminsearch, similarly to Ref. (47). This fitting is described in detail in *SI Appendix*.

For numerical ablations, the value of *Σ* was increased from *Σ* = 0 in these fitted shapes, with boundary conditions as in the toy problems above.

### Analytical calculations

Details of the analytical calculations for and additional discussion of the three mechanical toy problems, and the scaling argument for the recoil timescale are given in *SI Appendix*.

## ACKNOWLEDGMENTS

The authors are very grateful to Raymond E. Goldstein, in whose research group their collaboration on *Volvox* inversion mechanics started, for many discussions, mentoring, and invariably helpful scientific advice. The authors also thank members of the Cambridge Advanced Imaging Centre, particularly Martin Lenz and Kevin O’Holleran, for their advice and support. The authors gratefully acknowledge funding from the Max Planck Society (P.A.H.), and the Wellcome Trust and the John Templeton Foundation (S.S.M.H.H.).

## Footnotes

∗ We did not fit our model beyond the onset of phialopore opening because that is associated with poorly understood cell rearrangements near the phialopore (47), the mechanics of which cannot be described by our elastic model.

