## Supplemental Material for "Cut it out: Out-of-plane stresses in cell sheet folding of *Volvox* embryos"

### Supporting Information Text

This Supplementary Text is divided into four parts. The first three parts provide detailed calculations for the "toy problems" in the main text: We first analyse the (small) deformations of a spherical elastic shell with a homogeneous curvature mismatch. We then discuss the (larger) deformations resulting from a circular cut in such a shell, developing the theory and scaling arguments for laser ablations in curved tissues. We then discuss cell sheet curling due to inhomogeneous contraction rather than curvature mismatch. The final part describes the fitting of the elastic model to the averaged inversion shapes in detail.

### Toy Problem 1: deformations of a spherical elastic shell with mismatched intrinsic curvatures

As in the main text, we consider a thin incompressible elastic shell of undeformed (nondimensional) unit radius, and thickness $h \ll 1$. The shell has uniform intrinsic curvatures $\kappa^0 = k$. If $k \neq 1$, these are different from the curvatures of the undeformed shell. Due to this curvature mismatch, the shell deforms. We assume the shell to remain spherical and that $k = O(1)$, and denote by $f$ its deformed radius.

**Geometry of the deformed shell.** The (spherical) polar radius of a point in the undeformed configuration of the shell is $r = a + \zeta$, where $a$ is the midsurface radius, to be determined in the calculation, and $\zeta$ is a transverse coordinate, with $-h^- \leqslant \zeta \leqslant h^+$. By definition, $h^+ + h^- = h$, and $[(a + h^+) + (a - h^-)]/2 = 1$, i.e. $a = 1 - (h^+ - h^-)/2$. Analogously, a point in the deformed configuration of the shell has polar radius $\tilde{r} = \tilde{a} + \tilde{\zeta}$, where $\tilde{a}$ and $\tilde{\zeta}$ are the deformed midsurface radius and transverse coordinate, now with $-\tilde{h}^- \leqslant \tilde{\zeta} \leqslant \tilde{h}^+$ and hence $\tilde{a} = f - (\tilde{h}^+ - \tilde{h}^-)/2$. With respect to spherical polars $(r, \theta, \phi)$, the deformation gradient is diagonal, with principal stretches

$$\lambda_r = \frac{\partial \tilde{\zeta}}{\partial \zeta}, \qquad \lambda_\theta = \lambda_\phi = \frac{\tilde{a}}{a}\frac{1 + \tilde{\zeta}/\tilde{a}}{1 + \zeta/a}. \tag{S1}$$

Incompressibility requires $\lambda_r \lambda_\theta \lambda_\phi = 1$, which is a differential equation for $\tilde{\zeta}$ as a function of $\zeta$. Imposing $\tilde{\zeta} = 0$ at $\zeta = 0$, required by the definition of the midsurface, we find its implicit solution

$$\left(\frac{\tilde{a}}{a}\right)^2 \left(\tilde{\zeta} + \frac{\tilde{\zeta}^2}{\tilde{a}} + \frac{\tilde{\zeta}^3}{3\tilde{a}^2}\right) = \zeta + \frac{\zeta^2}{a} + \frac{\zeta^3}{3a^2}. \tag{S2}$$

We now make use of the framework of morphoelasticity (S1): By analogy with Eqs. **S1**, we define the intrinsic configuration of the cell sheet (which need not be embeddable into three-dimensional space) by its intrinsic stretches

$$\lambda_r^0 = \frac{\partial Z}{\partial \zeta}, \qquad \lambda_\theta^0 = \lambda_\phi^0 = \frac{1 + kZ}{1 + \zeta/a}. \tag{S3}$$

To write down these expressions, we have identified $\tilde{a}/a$ and $1/\tilde{a}$ in Eqs. **S1** as the stretch and curvature of the deformed configuration, respectively, and have replaced them with their respective intrinsic values 1 and $k$. We will refer to $Z$ as the intrinsic transverse coordinate. We are left to define the midsurfaces (a different choice of these would change the interpretation of $k$, so the existence of this freedom of choice is not surprising). We do so by imposing $-h^0/2 \leqslant Z \leqslant h^0/2$; the intrinsic thickness $h^0$ is determined by the incompressibility condition $\lambda_r^0 \lambda_\theta^0 \lambda_\phi^0 = 1$, which integrates to

$$\zeta + \frac{\zeta^2}{a} + \frac{\zeta^3}{3a^2} = Z + kZ^2 + \frac{k^2 Z^3}{3} \tag{S4}$$

on requiring $Z = 0$ at $\zeta = 0$. Imposing $Z = \pm h^0/2 \iff \zeta = \pm h^\pm \iff \tilde{\zeta} = \pm \tilde{h}^\pm$ in Eqs. **S2** and **S4** yields

$$\left(\frac{f - \dfrac{\tilde{h}^+ - \tilde{h}^-}{2}}{1 - \dfrac{h^+ - h^-}{2}}\right)^2 \left[\pm\tilde{h}^\pm + \frac{\left(\tilde{h}^\pm\right)^2}{f - \dfrac{\tilde{h}^+ - \tilde{h}^-}{2}} \pm \frac{\left(\tilde{h}^\pm\right)^3}{3\left(f - \dfrac{\tilde{h}^+ - \tilde{h}^-}{2}\right)^2}\right] = \pm h^\pm + \frac{\left(h^\pm\right)^2}{1 - \dfrac{h^+ - h^-}{2}} \pm \frac{\left(h^\pm\right)^3}{3\left(1 - \dfrac{h^+ - h^-}{2}\right)^2} \tag{S5a}$$

$$= \pm\frac{h^0}{2} + \frac{k\left(h^0\right)^2}{4} \pm \frac{k^2\left(h^0\right)^3}{24}. \tag{S5b}$$

Together with the condition $h^+ + h^- = h$, these form a system of equations for $h^\pm$, $\tilde{h}^\pm$, and $h^0$, the solution of which defines the deformed geometry of the shell.

**Elasticity of the deformed shell.** Let $\boldsymbol{F}$ and $\boldsymbol{F^0}$ be the diagonal tensors with principal stretches $\lambda_r, \lambda_\theta, \lambda_\phi$ and $\lambda_r^0, \lambda_\theta^0, \lambda_\phi^0$, respectively, relative to the standard basis of spherical polars. By the fundamental relation of morphoelasticity (S1), the elastic deformation gradient is $\boldsymbol{F}\left(\boldsymbol{F^0}\right)^{-1}$. The elastic deformation gradient therefore has principal stretches

$$\Lambda_\theta = \Lambda_\phi = \frac{\lambda_\theta}{\lambda_\theta^0} = \frac{\lambda_\phi}{\lambda_\phi^0} = \frac{\tilde{a}}{a}\frac{1 + \tilde{\zeta}/\tilde{a}}{1 + kZ} \equiv \Lambda, \qquad \Lambda_r = \frac{\lambda_r}{\lambda_r^0} = \frac{1/\lambda_\theta\lambda_\phi}{1/\lambda_\theta^0\lambda_\phi^0} = \frac{1}{\Lambda^2}, \tag{S6}$$

where we have used the incompressibility condition. The first two principal invariants of the associated Cauchy–Green tensor are thus $\mathcal{I}_1 = 2\Lambda^2 + \Lambda^{-4}$ and $\mathcal{I}_2 = \Lambda^4 + 2\Lambda^{-2}$.

Since the problem that we are solving is *not* a leading-order problem (in an expansion in the small thickness of the shell), the constitutive assumptions matter. We therefore consider general elastic energy densities with a regular power series expansion (S2), viz.,

$$e = \frac{1}{2}\sum_{m=0}^{\infty}\sum_{n=0}^{\infty} C_{mn}(\mathcal{I}_1 - 3)^m(\mathcal{I}_2 - 3)^n, \tag{S7}$$

where we have set $C_{00} = 0$ without loss of generality, and we may assume that $C_{01} + C_{10} > 0$ for the material to have a positive bulk modulus (S3). By incompressibility, we have equality $\mathrm{d}V^0 = \mathrm{d}V$ of the volume elements of the intrinsic and undeformed configurations. Explicitly,

$$\mathrm{d}V^0 = \mathrm{d}V = (a+\zeta)^2 \sin\theta\,\mathrm{d}\zeta\,\mathrm{d}\theta\,\mathrm{d}\phi = ha^2(1+khz)^2\sin\theta\,\mathrm{d}z\,\mathrm{d}\theta\,\mathrm{d}\phi, \tag{S8}$$

using incompressibility again, and where $z = Z/h$. The elastic energy of the shell is the integral of the energy density with respect to the intrinsic configuration, viz.,

$$\mathcal{E} = \int_{\mathcal{V}^0} e\,\mathrm{d}V^0 = 4\pi h\int_{-h^0/2h}^{h^0/2h} ea^2(1+khz)^2\,\mathrm{d}z. \tag{S9}$$

**Asymptotic solution.** We now determine the deformed radius $f$ by minimising $\mathcal{E}$ by asymptotic expansion for $h \ll 1$. To keep the algebraic expressions that arise in the calculation (somewhat) simple, it will be useful to divide the asymptotic solution into two steps.

***Leading-order calculation.*** We begin by showing that $f = 1 + O(h^2)$, recovering the scaling behaviour obtained in the main text using a shell theory. To this end, we introduce expansions

$$h^{\pm} = \frac{h}{2} + h^2h_2^{\pm} + O(h^3), \quad \tilde{h}^{\pm} = \frac{h}{2} + h^2\tilde{h}_2^{\pm} + O(h^3), \quad h^0 = h + h^2h_2^0 + O(h^3), \quad f = 1 + hf_1 + h^2f_2 + O(h^3). \tag{S10}$$

Using Mathematica (Wolfram, Inc.), Eqs. **S5** yield

$$h_2^+ = -h_2^- = \frac{k-1}{4}, \qquad \tilde{h}_2^+ = \frac{k-1}{4} - f_1, \qquad \tilde{h}_2^- = -\frac{k-1}{4} - f_1, \qquad h_2^0 = 0. \tag{S11}$$

We then set $Z = hz$, $\tilde{\zeta} = h\tilde{\zeta}_1 + h^2\tilde{\zeta}_2 + O(h^3)$ and solve Eqs. **S2** and **S4** for $\tilde{\zeta}_1, \tilde{\zeta}_2$ as functions of $z$. We find

$$\tilde{\zeta}_1 = z, \qquad \tilde{\zeta}_2 = -2f_1z + (k-1)z^2, \tag{S12}$$

and hence compute

$$\mathcal{E} = 2\pi(C_{10} + C_{01})\bigl[12f_1^2 + (k-1)^2\bigr]h^2 + O(h^3), \tag{S13}$$

which is minimised for all $k$ at $f_1 = 0$. This proves that $f = 1 + O(h^2)$ as claimed.

***Next-order calculation.*** To determine the leading-order deformation of the shell, we extend expansions **S10** using the leading-order results **S11**, writing

$$h^{\pm} = \frac{h}{2} \pm \frac{h^2(k-1)}{4} + h^3h_3^{\pm} + h^4h_4^{\pm} + O(h^5), \qquad \tilde{h}^{\pm} = \frac{h}{2} \pm \frac{h^2(k-1)}{4} + h^3\tilde{h}_3^{\pm} + h^4\tilde{h}_4^{\pm} + O(h^5), \tag{S14a}$$

and

$$h^0 = h + h^3h_3^0 + h^4h_4^0 + O(h^5), \qquad f = 1 + h^2f_2 + h^3f_3 + h^4f_4 + O(h^5). \tag{S14b}$$

Solving Eqs. **S5** order-by-order for $h_3^{\pm}, h_4^{\pm}, \tilde{h}_3^{\pm}, \tilde{h}_4^{\pm}, h_3^0, h_4^0$, leaving only $f_2, f_3, f_4$ as unknowns, we find

$$h_3^{\pm} = 0, \qquad \tilde{h}_3^{\pm} = -f_2, \qquad h_3^0 = -\frac{1}{12}\left(5 - 6k + k^2\right) \tag{S15a}$$

and

$$h_4^{\pm} = \pm\frac{1}{48}\left(3 - 10k + 9k^2 - 2k^3\right), \qquad \tilde{h}_4^{\pm} = \pm\frac{1}{48}\left(3 - 10k + 9k^2 - 2k^3\right) - f_3 \pm \frac{f_2}{4}(5-2k), \qquad h_4^0 = 0. \tag{S15b}$$

We set $Z = hz$ again and write $\tilde{\zeta} = hz + (k-1)h^2z^2 + \tilde{\zeta}_3h^3 + \tilde{\zeta}_4h^4 + O(h^5)$ using the leading-order results **S12**. From Eqs. **S2** and **S4**, we obtain

$$\tilde{\zeta}_3 = -2f_2z + \frac{z^3}{3}\left(5 - 6k + k^2\right), \qquad \tilde{\zeta}_4 = -2f_3z - \frac{z^2}{4}\left[k - 1 - 4(5-2k)f_2\right] - \frac{z^4}{3}\left(10 - 15k + 9k^2\right). \tag{S16}$$

Using MATHEMATICA, we compute

$$\frac{\mathcal{E}}{2\pi} = (C_{10}+C_{01})(k-1)^2h^2 + \Bigl\{12(C_{10}+C_{01})f_2^2 - (k-1)[C_{10}(5k-11)+C_{01}(k-7)]f_2 + g(k;C_{10},C_{01},C_{20},C_{11},C_{02})\Bigr\}\,h^4 + O(h^5), \qquad \textbf{[S17]}$$

where $g$ is a quartic polynomial in $k$ with coefficients depending on the material parameters $C_{10}, C_{01}, C_{20}, C_{11}, C_{02}$, the explicit expression of which is of no relevance to our calculation. Minimising $\mathcal{E}$ to determine $f_2$ finally yields

$$f = 1 - \frac{h^2}{12}(k-1)\frac{(11C_{10}+7C_{01})-(5C_{10}+C_{01})k}{2(C_{10}+C_{01})} + O(h^3). \qquad \textbf{[S18]}$$

***Surprising mechanical behaviour.*** While this result has the same scaling behaviour as the result derived in the main text from a shell theory not formally valid for these small deformations, it features a complicated prefactor depending on the material parameters $C_{10}, C_{01}$ and also on $k$ that is not predicted by the shell theory, but that gives rise to surprising and somewhat counterintuitive behaviour: indeed, the $O(h^2)$ correction in Eq. **S18** vanishes and changes sign not only at $k = 1$ as expected, but also at $k = k_*$, where $k_* = (11C_{10}+7C_{01})/(5C_{10}+C_{01}) = 1+6(C_{10}+C_{01})/(5C_{10}+C_{01})$. From $C_{10}+C_{01} > 0$, as required for the material to have a positive bulk modulus, it follows that $k_* \gtrless 1$ iff $5C_{10}+C_{01} \gtrless 0$, and hence $f \gtrless 1$ iff $1 \lessgtr k \lessgtr k_*$ for $5C_{10}+C_{01} \gtrless 0$. As expected, this shows that the sphere grows (shrinks) if $k$ is just below (above) unit value, but it is surprising that the sphere grows if $k > k_*$ ($> 1$) if $5C_{10}+C_{01} > 0$ and shrinks if $k < k_*$ ($< 1$) if $5C_{10}+C_{01} < 0$. It is likely that the reason for which the intuition that the sphere should simply grow and shrink if $k < 1$ and $k > 1$ respectively fails because these are very small deformations beyond leading order, asymptotically smaller than the thickness of the shell.

In this context, we also note that (again because the problem is not a leading-order problem), the definition of the midsurfaces (and hence of the intrinsic curvatures) matters: Had we chosen to define the midsurface in the undeformed configuration rather than in the intrinsic configuration, we would have obtained a different answer, simply because the middle surface of the undeformed configuration does not map to the middle surface of the intrinsic configuration, and so the intrinsic curvatures do not directly correspond to each other. This ambivalence does not however change the fact that there is "surprising" behaviour as discussed in the previous paragraph.

### Toy Problem 2: circular ablation of a spherical elastic shell with mismatched intrinsic curvatures

As in the main text, we consider a circular cut of angular extent $2\Sigma$ in a thin incompressible elastic spherical shell of unit radius, of thickness $h \ll 1$, and with intrinsic curvatures $\kappa^0 = k$. If $k \neq 1$, these are mismatched; to determine the resulting deformation of the shell, we take cylindrical polar coordinates defined by the axis of the cut.

**Governing equations and boundary conditions.** With respect to these coordinates, the undeformed spherical shell has polar radius $\rho(s) = \sin s$, where $s$ is arclength, measured from the centre of the cut, so its rim corresponds to $s = \Sigma$. The deformed shell has arclength $S(s)$, corresponding polar radius $r(s)$, and tangent angle $\psi(s)$. The stretches and curvatures of the deformed shell are thus

$$f_s(s) = \frac{\mathrm{d}S}{\mathrm{d}s}, \qquad f_\phi(s) = \frac{r(s)}{\rho(s)}, \qquad \kappa_s(s) = \frac{1}{f_s}\frac{\mathrm{d}\psi}{\mathrm{d}s}, \qquad \kappa_\phi(s) = \frac{\sin\psi(s)}{r(s)}. \qquad \textbf{[S19]}$$

These define the shell strains and curvature strains

$$E_s = f_s - 1, \qquad E_\phi = f_\phi - 1, \qquad K_s = f_s\kappa_s - k, \qquad K_\phi = f_\phi\kappa_\phi - k. \qquad \textbf{[S20]}$$

For this toy problem, the curvatures remain small compared to the shell thickness. We do not therefore need to use the "large bending theory" (S3) used in the numerical calculations in the main text for our asymptotic calculations. On defining

$$N_s = 2E_s + E_\phi, \qquad N_\phi = E_s + 2E_\phi, \qquad M_s = 2K_s + K_\phi, \qquad M_\phi = K_s + 2K_\phi, \qquad \textbf{[S21]}$$

and letting $\varepsilon = h/\sqrt{12}$, the deformed configuration of the shell satisfies the (nondimensionalised) force-balance equations (S4)

$$\varepsilon^2\frac{\mathrm{d}}{\mathrm{d}s}(rM_s) = f_s\left(rN_s\tan\psi + \varepsilon^2 M_\phi\cos\psi\right), \qquad \frac{\mathrm{d}}{\mathrm{d}s}(rN_s\sec\psi) = f_sN_\phi, \qquad \textbf{[S22a]}$$

complemented by the geometric equations

$$\frac{\mathrm{d}r}{\mathrm{d}s} = f_s\cos\psi, \qquad \frac{\mathrm{d}\psi}{\mathrm{d}s} = f_s\kappa_s. \qquad \textbf{[S22b]}$$

It will however turn out to be convenient to express these equations in terms of $\psi$, $E_s$, $E_\phi$, $N_s$ for the asymptotic solution. To this end, rearranging the first of Eqs. **S22b** using definitions **S20**, we notice that

$$\frac{\mathrm{d}}{\mathrm{d}s}\bigl(E_\phi\rho(s)\bigr) = f_s\cos\psi - \rho'(s). \qquad \textbf{[S23]}$$

Moreover, we obtain, from Eqs. **S20** and **S21**, the algebraic relations

$$2N_\phi = 3E_\phi + N_s, \qquad\qquad 2f_s = 2 + N_s - E_\phi. \tag{S24}$$

Inserting these into Eqs. **S22** and **S23** and expanding, we conclude that deformed configuration of the shell is described by the equations

$$4\frac{\mathrm{d}}{\mathrm{d}s}\left[(1+E_\phi)N_s\sec\psi\sin s\right] = (2+N_s-E_\phi)(3E_\phi+N_s), \tag{S25a}$$

$$2\varepsilon^2\frac{\mathrm{d}}{\mathrm{d}s}\left[(1+E_\phi)\sin s\left(2\frac{\mathrm{d}\psi}{\mathrm{d}s}+\frac{\sin\psi}{\sin s}-3k\right)\right] = (2+N_s-E_\phi)\left[(1+E_\phi)N_s\tan\psi\sin s+\varepsilon^2\cos\psi\left(\frac{\mathrm{d}\psi}{\mathrm{d}s}+2\frac{\sin\psi}{\sin s}-3k\right)\right], \tag{S25b}$$

$$2\frac{\mathrm{d}}{\mathrm{d}s}\left(E_\phi\sin s\right) = 2(\cos\psi-\cos s)+(N_s-E_\phi)\cos\psi. \tag{S25c}$$

The boundary conditions at the rim of the cut, $s=\varSigma$, are

$$N_s(\varSigma)=0, \qquad\qquad M_s(\varSigma)=0 \quad\Longleftrightarrow\quad 2\psi'(\varSigma)+\frac{\sin\psi(\varSigma)}{\sin\varSigma}=3k. \tag{S26}$$

**Asymptotic solution.** In the asymptotic limit $\varepsilon \ll 1$, the shell deforms in a region of characteristic extent $\delta \ll 1$ near $s=\varSigma$. In this inner region, we let $s=\varSigma+\delta\xi$, defining an inner coordinate $\xi$. Let $\beta(\xi)=\psi(s)-s$ be the deviation of the tangent angle due to the cut; this deviation is small, $\beta\ll 1$. Expanding the no-torque boundary condition,

$$\frac{2}{\delta}\dot{\beta}(0)+\beta(0)\cot\varSigma = 3(k-1), \tag{S27}$$

where the dot denotes a derivative with respect to $\xi$. This condition therefore requires $\beta=O(\delta)$. Scaling from Eqs. **S25**, we also find

$$\frac{N_s}{\delta}\sim E_\phi, \qquad\qquad \frac{\varepsilon^2\beta}{\delta^2}\sim N_s, \qquad\qquad \frac{E_\phi}{\delta}\sim\max\{\beta,N_s,E_\phi\}. \tag{S28}$$

The first scaling yields $N_s \ll E_\phi$, and hence the third scaling becomes $E_\phi/\delta\sim\beta$. The scalings then reduce to $\delta\sim\sqrt{\varepsilon}$, and hence $E_\phi\sim\delta^2$, $N_s\sim\delta^3$. [The asymptotic expansion of elasticity (S3) that leads to the shell equations **S22** includes the scalings $E_s, E_\phi = O(\varepsilon)$ and $K_s, K_\phi = O(1)$, consistent with these scalings.] We therefore posit regular expansions

$$\psi = s+\delta\left[b+O(\delta)\right], \qquad\qquad E_\phi=\delta^2\left[e+O(\delta)\right], \qquad\qquad N_s=\delta^3\cot\varSigma\left[n+O(\delta)\right], \tag{S29}$$

so that, on Taylor expanding and at leading order, Eqs. **S25** become

$$2\dot{n}=3e, \qquad\qquad 2\ddot{b}=n, \qquad\qquad \dot{e}=-b. \tag{S30}$$

The leading-order problem is therefore, on recalling Eqs. **S26** and **S27**,

$$4\ddddot{b}+3b=0 \qquad \text{subject to}\quad b\to 0 \text{ as } \xi\to\infty,\ 2\dot{b}(0)=3(k-1),\ \ddot{b}(0)=0. \tag{S31}$$

Its solution is

$$b(\xi) = -3^{3/4}(k-1)\mathrm{e}^{-3^{1/4}\xi/2}\cos\left(\frac{3^{1/4}\xi}{2}\right). \tag{S32}$$

***Cut opening.*** One experimentally accessible geometric measure that characterises the deformation resulting from the curvature mismatch is the cut opening $o$, i.e. the amount by which the cut opens during the recoil following the ablation. To compute $o$, we note that the radial displacement $u(s)=r(s)-\rho(s)$ of the shell satisfies the differential equation

$$\frac{\mathrm{d}u}{\mathrm{d}s}=f_s\cos\psi-\cos s \quad\Longrightarrow\quad \frac{\mathrm{d}u}{\mathrm{d}\xi}=-\delta^2 b\sin\varSigma+O(\delta^3), \tag{S33}$$

were we have made use of $f_s=1+O(\delta^2)$. Now $o=2[u(\xi=0)-u(\xi\to\infty)]$, twice the radial displacement of the cut edge across the boundary layer. We thus obtain

$$o=2\delta^2\sin\varSigma\int_0^\infty b(\xi)\,\mathrm{d}\xi+O(\delta^3) = -h(k-1)\sin\varSigma+O(\delta^3). \tag{S34}$$

As expected, this shows that the rim of the cut curls outwards if $k<1$, and inwards if $k>1$. We also note that $o$ is proportional to the radius $\sin\varSigma$ of the ablation.

***Cut rotation and cut displacement: breakdown of the leading-order asymptotics.*** There are (at least) two other exerimentally accessible measures of the deformation: the rotation $\varrho$ of the edge of the cut, and its displacement $d$ during the recoil. By definition,

$$\varrho = \delta b(0) + O(\delta^2) = -\sqrt{\frac{3h}{2}}(k-1) + O(\delta^2). \qquad \textbf{[S35]}$$

Moreover, the calculation of the cut opening shows that the radial displacement of the cut edge is $-(h/2)(k-1)\sin\Sigma + O(\delta^3)$. Similarly, the axial displacement of the cut edge is $-(h/2)(k-1)\cos\Sigma + O(\delta^3)$, whence

$$d = \frac{h}{2}|k-1| + O(\delta^3). \qquad \textbf{[S36]}$$

Both leading-order results are independent of the cut size $\Sigma$. This is clearly absurd, because we must have $\varrho, d \to 0$ as $\Sigma \to 0$. This signals a breakdown of the leading-order asymptotics that we will analyse in the next subsection.

**Asymptotic solution for small ablations.** We now address the breakdown of the leading-order asymptotic solution for small cut sizes that we highlighted above when deriving the leading-order expressions for the cut rotation and cut displacement.

***Breakdown of asymptoticity.*** To understand this breakdown of the asymptotic validity of the leading-order solution, we expand further, writing

$$\psi = s + \delta\left[b_0 + \delta b_1 + O(\delta^2)\right], \qquad E_\phi = \delta^2\left[e_0 + \delta e_1 + O(\delta^2)\right], \qquad N_s = \delta^3\cot\Sigma\left[n_0 + \delta n_1 + O(\delta^2)\right], \qquad \textbf{[S37]}$$

where $b_0$ is given by Eq. **S32**, and $n_0 = 2\ddot{b}_0$, $e_0 = 4\dddot{b}_0/3$ from Eqs. **S30**. On expanding Eqs. **S25** and the boundary conditions pertaining thereto using MATHEMATICA, we obtain

$$4\dddot{b}_1 + 3b_1 = \cot\Sigma\left(4\ddot{b}_0^2 + 8\dot{b}_0\dddot{b}_0 - 8\ddot{b}_0 - \tfrac{9}{2}b_0^2\right) \quad \text{subject to } \dot{b}_1(0) = -\tfrac{1}{2}b_0(0)\cot\Sigma,\ \ddot{b}_1(0) = -\dot{b}_0(0)\cot\Sigma. \qquad \textbf{[S38]}$$

This shows that $b_1(\xi)$ is proportional to $\cot\Sigma$, so the expansion $b = b_0 + \delta b_1 + O(\delta^2)$ loses asymptoticity when $\delta\cot\Sigma = O(1)$, which is when $\Sigma = O(\delta)$.

***Asymptotic scalings for small ablations.*** We must therefore derive the asymptotic scalings for small cut sizes $\Sigma = O(\delta)$. We thus write $\Sigma = \delta\sigma$, where $\sigma = O(1)$. The scaling $\beta = O(\delta)$ still holds, but establishing the other scalings takes more effort.

First, expanding Eq. **S25a** shows that $E_\phi \lesssim N_s$. If $E_\phi \ll N_s$, then we may posit $N_s = \nu n + o(\nu)$, $E_\phi = o(\nu)$ for some $\nu \ll 1$, and Eq. **S25a** becomes, at leading order,

$$2\frac{\mathrm{d}}{\mathrm{d}\xi}\left[(\sigma+\xi)n\right] = n \quad\Longrightarrow\quad n(\xi) = \frac{C_1}{\sqrt{\xi+\sigma}}, \qquad \textbf{[S39]}$$

where $C_1$ is a constant of integration. The no-force condition at the rim of the cut is $n(0) = 0$, yielding $C_1 = 0$ and hence $n \equiv 0$, which is a contradiction. This shows that $N_s \sim E_\phi$.

Next, on expanding Eq. **S25c**, we find that $\delta^2 \lesssim E_\phi \sim N_s$. If $\delta^2 \ll E_\phi \sim N_s$, then $N_s = \nu n + o(\nu)$, $E_\phi = \nu e + o(\nu)$, for some $\nu \ll 1$, and hence, from Eqs. **S25a** and **S25c**,

$$2\frac{\mathrm{d}}{\mathrm{d}\xi}\left[(\sigma+\xi)n\right] = 3e + n,\ 2\frac{\mathrm{d}}{\mathrm{d}\xi}\left[(\sigma+\xi)e\right] = n - e \quad\Longrightarrow\quad 2(\sigma+\xi)\dot{n} = 3e - n,\ 2(\sigma+\xi)\dot{e} = n - 3e. \qquad \textbf{[S40a]}$$

Summing these equations yields $n + e = \text{const.}$, and then integrating the first gives, in particular,

$$n(\xi) = C_2 + \frac{C_3}{(\sigma+\xi)^2}, \qquad \textbf{[S40b]}$$

wherein $C_2, C_3$ are more constants of integration. The no-force condition at the rim of the cut is $n(0) = 0$, while matching to the undeformed configuration $f_s = f_\phi = 1$ as $\xi \to \infty$ requires $n \to 0$ in this limit. Hence $C_2 = C_3 = 0$, so $n \equiv 0$. This is another contradiction, so $E_\phi \sim N_s \sim \delta^2$.

Finally, expanding Eq. **S25b** gives $N_s\delta^2 \lesssim \varepsilon^2$. If $N_s\delta^2 \ll \varepsilon^2$, then, on positing $\psi = s + \delta b + O(\delta^2)$ again, Eq. **S25b** becomes, at leading order,

$$\frac{\mathrm{d}}{\mathrm{d}s}\left[2(\sigma+\xi)\dot{b} + b\right] = \dot{b} + \frac{2b}{\sigma+\xi} \quad\Longrightarrow\quad (\xi+\sigma)^2\ddot{b} + (\xi+\sigma)\dot{b} - b = 0, \qquad \textbf{[S41a]}$$

which is a homogeneous equation with general solution

$$b(\xi) = C_4(\xi+\sigma) + \frac{C_5}{\xi+\sigma}, \qquad \textbf{[S41b]}$$

where $C_4, C_5$ are yet more constants of integrations. Matching to the undeformed shell as $\xi \to \infty$ requires $\beta \to 0$ in this limit, and hence $C_4 = 0$. Let $v(s)$ be the vertical displacement of the shell, which satisfies $\mathrm{d}v/\mathrm{d}s = f_s\sin\psi - \sin s$. Expanding yields $\dot{v} = \delta^2 C_5(\xi+\sigma)^{-1} + O(\delta^3)$, which does not integrate to a finite displacement unless $C_5 = 0$, i.e. unless $b \equiv 0$.

This is a final contradiction, so $N_s\delta^2 \sim \varepsilon^2$. Since we have shown above that $E_\phi \sim N_s \sim \delta^2$, this yields $\delta \sim \sqrt{\varepsilon}$, which finally determines the asymptotic balance. In particular, $E_s, E_\phi = O(\varepsilon)$, as expected from the scalings of the asymptotic expansion underlying the derivation of the shell theory (S3).

***Asymptotic governing equations for small ablations.*** Having determined the asymptotic balance for $\varSigma = O(\delta)$, we can now obtain the governing equations in this limit. According to the scalings derived above, we replace Eqs. **S29** with

$$\psi = s + \delta\left[b + O(\delta)\right], \qquad E_\phi = \delta^2\left[e + O(\delta)\right], \qquad N_s = \delta^2\left[n + O(\delta)\right]. \tag{S42}$$

Equations **S25** yield

$$2(\sigma+\xi)\dot{n} = 3e - n, \quad 2(\sigma+\xi)^2\ddot{b} = -2(\sigma+\xi)\dot{b} + 2b + (\sigma+\xi)^3 n + (\sigma+\xi)^2 bn, \quad 2(\sigma+\xi)\dot{e} = n - 3e - 2(\sigma+\xi)b - b^2. \tag{S43}$$

On eliminating $e$, these equations reduce to the pair of nonlinear second-order equations

$$4(\sigma+\xi)^2\ddot{n} = -12(\sigma+\xi)\dot{n} - 6(\sigma+\xi)b - 3b^2, \qquad 2(\sigma+\xi)^2\ddot{b} = -2(\sigma+\xi)\dot{b} + 2b + (\sigma+\xi)^3 n + (\sigma+\xi)^2 bn, \tag{S44}$$

which are subject to the boundary conditions

$$n(0) = 0, \qquad 2\dot{b}(0) + \frac{b(0)}{\sigma} = 3(k-1), \qquad b, n \to 0 \text{ as } \xi \to \infty. \tag{S45}$$

Since Eqs. **S44** are nonlinear, their solution is not proportional to $k-1$, unlike the solution of the leading-order problem for $\varSigma = O(1)$. They must be solved numerically in general.

***Approximate solution for small ablations.*** It is therefore all the more remarkable that it is possible to find an analytical approximate solution of Eqs. **S44** that satisfies the boundary conditions **S45** and for which the cut rotation and cut displacement match onto Eqs. **S35** and **S36** in the limit $\sigma \to \infty$.

Our starting point is the observation that the boundary conditions for $\xi \to \infty$ imply that $b \ll \sigma + \xi$ there, and hence the nonlinear terms in Eqs. **S44** can be neglected there. On setting $x = \sigma + \xi$, Eqs. **S44** reduce to

$$2x^2\ddot{n} = -6x\dot{n} - 3xb, \qquad 2x^2\ddot{b} = -2x\dot{b} + 2b + x^3 n, \tag{S46}$$

in the limit $x \to \infty$, where dots now denote derivatives with respect to $x$. The solution of these equations subject to boundary conditions **S45** will constitute our approximate solution.

To obtain this solution, we introduce the differential operator $\mathscr{D} = x^2\mathrm{d}^2/\mathrm{d}x^2 + x\mathrm{d}/\mathrm{d}x - 1$. By definition, $\mathscr{D}b = x^2(xn)/2$. Observe that $\mathscr{D}(xn) = x^3\ddot{n} + 3x^2\dot{n} = -3x^2 b/2$ using Eqs. **S46**, so

$$\mathscr{D}\left(b \pm \frac{\mathrm{i}}{\sqrt{3}}xn\right) = \mp\frac{\sqrt{3}}{2}\mathrm{i}x^2\left(b \pm \frac{\mathrm{i}}{\sqrt{3}}xn\right) = \left(\frac{3^{1/4}}{\sqrt{2}}\mathrm{e}^{\mp\mathrm{i}\pi/4}x\right)^2\left(b \pm \frac{\mathrm{i}}{\sqrt{3}}xn\right), \tag{S47}$$

which are complex Bessel equations (S5), the solutions of which can be written as

$$b + \frac{\mathrm{i}}{\sqrt{3}}xn = c_1\mathcal{K}_1\left(\frac{3^{1/4}}{\sqrt{2}}\mathrm{e}^{-\mathrm{i}\pi/4}x\right) + c_2\mathcal{I}_1\left(\frac{3^{1/4}}{\sqrt{2}}\mathrm{e}^{-\mathrm{i}\pi/4}x\right), \quad b - \frac{\mathrm{i}}{\sqrt{3}}xn = c_3\mathcal{K}_1\left(\frac{3^{1/4}}{\sqrt{2}}\mathrm{e}^{\mathrm{i}\pi/4}x\right) + c_4\mathcal{I}_1\left(\frac{3^{1/4}}{\sqrt{2}}\mathrm{e}^{\mathrm{i}\pi/4}x\right), \tag{S48}$$

where $c_1, c_2, c_3, c_4$ are constants of integration, and where $\mathcal{I}_1$, $\mathcal{K}_1$ are the modified Bessel functions of the first order (S5) which satisfy $\mathcal{I}_1(z) \sim (2\pi z)^{-1/2}\mathrm{e}^{z}$ and $\mathcal{K}_1(z) \sim (\pi/2z)^{1/2}\mathrm{e}^{-z}$ as $z \to \infty$ for $\mathrm{Re}(z) > 0$. Accordingly, $b, n \to 0$ as $x \to \infty$ requires $c_2 = c_4 = 0$. Hence

$$b(x) = \frac{c_1}{2}\mathcal{K}_1\left(\frac{3^{1/4}}{\sqrt{2}}\mathrm{e}^{-\mathrm{i}\pi/4}x\right) + \frac{c_3}{2}\mathcal{K}_1\left(\frac{3^{1/4}}{\sqrt{2}}\mathrm{e}^{\mathrm{i}\pi/4}x\right), \quad n(x) = \frac{\sqrt{3}}{x}\left[-\frac{\mathrm{i}c_1}{2}\mathcal{K}_1\left(\frac{3^{1/4}}{\sqrt{2}}\mathrm{e}^{-\mathrm{i}\pi/4}x\right) + \frac{\mathrm{i}c_3}{2}\mathcal{K}_1\left(\frac{3^{1/4}}{\sqrt{2}}\mathrm{e}^{\mathrm{i}\pi/4}x\right)\right], \tag{S49a}$$

$$= c_1'\operatorname{ker}_1\left(\frac{3^{1/4}x}{\sqrt{2}}\right) + c_2'\operatorname{kei}_1\left(\frac{3^{1/4}x}{\sqrt{2}}\right), \qquad = \frac{\sqrt{3}}{x}\left[-c_1'\operatorname{kei}_1\left(\frac{3^{1/4}x}{\sqrt{2}}\right) + c_2'\operatorname{ker}_1\left(\frac{3^{1/4}x}{\sqrt{2}}\right)\right], \tag{S49b}$$

where $c_1' = \mathrm{i}(c_3 - c_1)/2$, $c_2' = -(c_1 + c_3)/2$, and where we use $\operatorname{ker}_0, \operatorname{ker}_1, \dots$ and $\operatorname{kei}_0, \operatorname{kei}_1, \dots$ to denote the Kelvin functions (S5). Applying the remaining, first two boundary conditions in Eqs. **S45** using Mathematica and setting $\varsigma = 3^{1/4}\sigma/\sqrt{2}$, we obtain

$$c_1' = \frac{3^{3/4}\sqrt{2}(1-k)\varsigma\operatorname{ker}_1\varsigma}{(\operatorname{ker}_1\varsigma)^2 + (\operatorname{kei}_1\varsigma)^2 + \sqrt{2}\varsigma\left[\operatorname{ker}_1\varsigma\left(\operatorname{ker}_0\varsigma + \operatorname{kei}_0\varsigma\right) + \operatorname{kei}_1\varsigma\left(\operatorname{kei}_0\varsigma - \operatorname{ker}_0\varsigma\right)\right]}, \tag{S50a}$$

$$c_2' = \frac{3^{3/4}\sqrt{2}(1-k)\varsigma\operatorname{kei}_1\varsigma}{(\operatorname{ker}_1\varsigma)^2 + (\operatorname{kei}_1\varsigma)^2 + \sqrt{2}\varsigma\left[\operatorname{ker}_1\varsigma\left(\operatorname{ker}_0\varsigma + \operatorname{kei}_0\varsigma\right) + \operatorname{kei}_1\varsigma\left(\operatorname{kei}_0\varsigma - \operatorname{ker}_0\varsigma\right)\right]}. \tag{S50b}$$

From this solution, we obtain the leading-order rotatation of the edge of the cut

$$\varrho \sim \delta b(\sigma) = \frac{\sqrt{3}\varsigma\left[(\operatorname{ker}_1\varsigma)^2 + (\operatorname{kei}_1\varsigma)^2\right]}{(\operatorname{ker}_1\varsigma)^2 + (\operatorname{kei}_1\varsigma)^2 + \sqrt{2}\varsigma\left[\operatorname{ker}_1\varsigma\left(\operatorname{ker}_0\varsigma + \operatorname{kei}_0\varsigma\right) + \operatorname{kei}_1\varsigma\left(\operatorname{kei}_0\varsigma - \operatorname{ker}_0\varsigma\right)\right]}\sqrt{h}(1-k). \tag{S51}$$

In particular, $\varrho \to 0$ as $\sigma \to 0$: the cut rotation vanishes with the cut size as expected, so this solution resolves the unphysical behaviour of the expression in Eq. **S35** computed from the leading-order solution. Remarkably, we find using MATHEMATICA that $\varrho \to \sqrt{3h/2}(1-k)$ as $\sigma \to \infty$, recovering the result for $\varSigma = O(1)$ obtained earlier.

This solution resolves the unphysical behaviour in Eq. **S36** for the cut displacement in a similar way: The leading-order equations for the radial and vertical displacements of the shell are now

$$\frac{\mathrm{d}u}{\mathrm{d}\xi} = -\delta^3\left[\frac{b^2}{2} + (\sigma+\xi)b - f\right], \qquad \frac{\mathrm{d}v}{\mathrm{d}\xi} = \delta^2 b, \qquad \textbf{[S52]}$$

where $f(\xi) = [n(\xi) - e(\xi)]/2 = [n(\xi) - (\sigma+\xi)\dot{n}(\xi)]/3$ using the first of Eqs. **S43**. The first equation cannot be integrated in closed form, so the approximate solution does not yield a correction to the expression for the cut opening in a straightforward manner. However, Eqs. **S52** show that $v \gg u$ for $\varSigma = O(\delta)$, whence the cut edge displacement is $d \sim |v(\xi=0) - v(\xi \to \infty)|$. This leads to

$$d = \frac{\varsigma\left[\ker_1\varsigma\,(\mathrm{kei}_0\,\varsigma - \ker_0\varsigma) - \mathrm{kei}_1\,\varsigma\,(\mathrm{kei}_0\,\varsigma + \ker_0\varsigma)\right]}{\sqrt{2}\left\{(\ker_1\varsigma)^2 + (\mathrm{kei}_1\,\varsigma)^2 + \sqrt{2}\varsigma\left[\ker_1\varsigma\,(\ker_0\varsigma + \mathrm{kei}_0\,\varsigma) + \mathrm{kei}_1\,\varsigma\,(\mathrm{kei}_0\,\varsigma - \ker_0\varsigma)\right]\right\}} h|1-k|. \qquad \textbf{[S53]}$$

We find $d \to h|1-k|/2$ as $\sigma \to \infty$, which recovers the result for $\varSigma = O(1)$. This is quite remarkable, because the radial displacement, which is negligible for $\varSigma = O(\delta)$ contributes to the expression for $d$ obtained for $\varSigma = O(1)$. Moreover, $d \to 0$ as $\sigma \to 0$, resolving the limitation of the solution for $\varSigma = O(1)$ as announced.

**Recoil velocity: scaling argument.** With the leading-order asymptotic solution, we can understand the scaling of the velocity of the recoil from the undeformed to the deformed solution. If the shell were to remain in its undeformed configuration,

$$E_s = E_\phi = 0, \qquad K_s = K_\phi = 1-k. \qquad \textbf{[S54]}$$

The deformed solution has $N_s \ll E_\phi$ from Eqs. **S29**, whence, from the first of Eqs. **S21**, $E_s = -E_\phi/2 + O(\delta)$. It follows that, in the asymptotic inner region,

$$E_s \sim -\tfrac{1}{2}\delta^2 e, \qquad E_\phi \sim \delta^2 e, \qquad \textbf{[S55a]}$$

so definitions **S20** imply that $f_s, f_\phi \sim 1$. Hence, from Eqs. **S20** and **S29**,

$$K_s \sim 1 - k + \dot{b}, \qquad K_\phi \sim 1-k. \qquad \textbf{[S55b]}$$

The elastic energy density (S4) is thus

$$E_s^2 + E_sE_\phi + E_\phi^2 + \varepsilon^2\left(K_s^2 + K_sK_\phi + K_\phi^2\right) \sim \begin{cases} 3\varepsilon^2(1-k)^2 & \text{in the undeformed configuration,} \\ \dfrac{3}{4}\delta^4 e^2 + \varepsilon^2\left[3(1-k)(1-k+\dot{b}) + \dot{b}^2\right] & \text{in the deformed configuration.}\end{cases} \qquad \textbf{[S56]}$$

On noting that $e = \frac{4}{3}\ddot{b}$ from Eqs. **S30**, it follows that the elastic energy released during the recoil from the undeformed to the deformed configuration is

$$\Delta\mathcal{E} = 2\pi\int_0^\infty \delta^4\left[\frac{4}{3}\ddot{b}^2 + \dot{b}^2 + 3\dot{b}(1-k)\right]\sin\varSigma\,(\delta\,\mathrm{d}\xi) = -3^{7/4}\pi\delta^5(k-1)^2\sin\varSigma \leqslant 0, \qquad \textbf{[S57]}$$

using the leading-order solution **S32** and in which the factor $\sin\varSigma$ is the leading-order Jacobian of polar coordinates. Of course, the fact that elastic energy is released is hardly surprising, but we needed to compute this number to find the leading-order scaling of this energy release.

We now make this result dimensional: in our derivation of the leading-order solution, we have implicitly nondimensionalised distances with the dimensional radius $R$ of the shell, and energies with $YR^2$, where $Y = EH$ is the Young's modulus of the shell, with $E$ its elastic modulus and $H$ is dimensional thickness, so that $h = H/R$. On unravelling this nondimensionalisation, the dimensional energy released during the recoil thus scales as $\Delta\mathcal{E} \sim EHR^2\delta^5(k-1)^2\sin\varSigma$. During the recoil, the sheet moves a (dimensional) distance $D \sim R\delta^2|1-k|\sin\varSigma \sim H|1-k|\sin\varSigma$ from Eq. **S36**. The effective elastic force driving the recoil is thus $F \sim \Delta\mathcal{E}/D \sim EHR\delta^3|1-k| \sim EH^{5/2}R^{-1/2}|1-k|$. The recoil is resisted by viscous dissipation in the tissue and the surrounding fluid. The hydrodynamic strain rate scales as $\mu V/(\delta R)$, where $\mu$ is an effective viscosity and $V$ is the initial recoil velocity, i.e. the fluid velocity that decays over the extent $\delta R$ of the asymptotic inner region. The later has surface area $R^2\delta$, and so $F \sim (\mu V/\delta R)(R^2\delta) \sim \mu VR$. The initial recoil velocity and hence the hydrodynamic force are independent, at leading order, of $k$ because the initial shape of the cell sheet is, by the results of toy problem 1. The force balance of elastic and hydrodynamic forces then yields the scaling of the recoil velocity

$$V \sim \frac{E}{\mu}\frac{H^{5/2}}{R^{3/2}}|1-k|. \qquad \textbf{[S58]}$$

This scaling breaks down for small cut sizes, because it does not satisfy $V \to 0$ as $\varSigma \to 0$, but resolving this would require a more detailed understanding of the dependence of the fluid velocity on cut size and its decay away from the rim of the cut, which is beyond the scope of our analysis.

### Toy Problem 3: circular ablation of a spherical elastic shell with anisotropic contraction

Finally, we show how (spatially uniform) anisotropic contraction of a spherical shell of unit radius leads to stresses (and hence to recoil on ablation) irrespective of the intrinsic curvatures of the shell. Let $f_1 \neq f_2$ be the principal contractions, and take cylindrical coordinates such that the (orthogonal) principal directions of contraction align with the azimuthal and circumferential directions of the shell. As in the previous section, we denote by $s$ and $S(s)$ the undeformed and deformed arclengths of the shell along its meridian (measured from a pole), and by $\rho(s) = \sin s$ and $r(s)$ the undeformed and deformed polar radii of the shell. The strains of the midsurface of the shell (S3) are thus

$$E_s(s) = \frac{f_s(s)}{f_1} - 1, \qquad E_\phi(s) = \frac{f_\phi(s)}{f_2} - 1, \qquad \text{where } f_s(s) = S'(s),\ f_\phi(s) = \frac{r(s)}{\rho(s)}, \qquad \textbf{[S59]}$$

and where dashes denote differentiation with respect to $s$. Let $\psi(s)$ denote the deformed tangent angle to the shell, which satisfies $r'(s) = f_s \sin\psi$. At the pole of the shell, we have $r(0) = \rho(0) = \psi(0) = 0$ by geometric continuity, so l'Hôpital's rule implies $f_\phi(0) = r'(0)/\rho'(0) = f_s(0)\cos\psi(0)/\cos 0 = f_s(0)$. Since $f_1 \neq f_2$, it follows that $E_s(0)$ and $E_\phi(0)$ cannot both vanish, so the meridional and circumferential stresses $N_s = 2E_s + E_\phi$ and $N_\phi = E_s + 2E_\phi$, the expressions of which are derived in Ref. (S4), cannot both vanish near the pole by continuity, and hence the shell will recoil on ablation.

While these stresses are caused by anisotropic contraction, we note that they also generate a curvature mismatch. Indeed, let $K_s, K_\phi$ denote the mismatch of the actual and intrinsic curvatures of the shell in the meridional and circumferential directions. Then $M_s = 2K_s + K_\phi$, $M_\phi = K_s + 2K_\phi$ satisfy the differential equation $\varepsilon^2(rM_s)' = f_s(rN_s\tan\psi + \varepsilon^2 M_\phi\cos\psi)$ as derived in Ref. (S4), where $\varepsilon$ is again the (scaled) relative thickness of the shell. Hence $M_s, M_\phi$ cannot both vanish identically since $N_s$ does not, and so neither can $K_s, K_\phi$ as claimed.

### Fitting of the morphoelastic theory to the average cell sheet midlines of *Volvox* inversion

We fit our morphoelastic model (*Materials and Methods* of the main text), based on the elastic shell theory for large bending deformations of Ref. (S3), to the average cell sheet midlines of *Volvox* inversion (S6) similarly to the fitting described in Ref. (S6). The use of the upgraded shell theory of Ref. (S3) is the main mechanical difference between the quantitative inversion model of this paper and that of Ref. (S6). In what follows, we describe the quantitative model and the fitting in detail.

**Fitting parameters encode cell shape changes in intrinsic stretches and curvatures.** Similarly to Ref. (S6), we define piecewise constant or linear functional forms for the intrinsic stretch and intrinsic curvature functions $f_s^0, f_\phi^0, \kappa_s^0, \kappa_\phi^0$ (*Materials and Methods* of the main text) that encode the cell shape changes observed during *Volvox* inversion (S7). These functional forms are shown in Fig. S1. They define fitting parameters $f_1, f_2, f_3, f_4, f_5$, $\kappa_1, \kappa_2, \kappa_3, \kappa_4$, $s_1 < s_2 < s_3 < s_4$ that encode the intrinsic stretches and intrinsic curvatures of the different cell shapes observed during *Volvox* inversion, and the positions of the boundaries between regions of different cell shapes. There are thus 13 fitting parameters. Since the intrinsic stretches and curvatures change during development, as the programme of cell shape driving inversion is run through, each of these parameters is *a priori* a function of time, so has to be fitted for at each developmental timepoint. This number of fitting parameters may therefore appear to be large, but we emphasise, as discussed in detail in Ref. (S6), that the functional forms of intrinsic stretches and curvatures are the minimal ones that can encode the cell shape changes observed during *Volvox* inversion (S7) and are appropriately continuous.

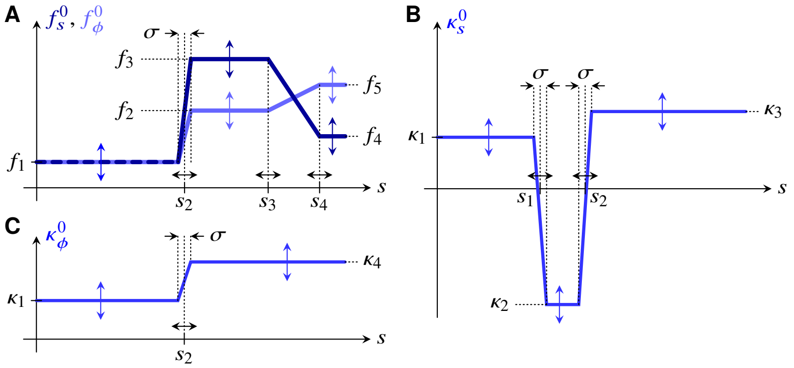


**Fig. S1.** Functional forms of intrinsic stretches and curvatures. (A) Plots of the piecewise linear or constant functional forms of the intrinsic stretches $f_s^0, f_\phi^0$ against arclength $s$ (*Materials and Methods* of the main text), defining fitting parameters $f_1, f_2, f_3, f_4, f_5$, $s_2 < s_3 < s_4$ encoding the intrinsic stretches of the different cell shapes observed during *Volvox* inversion (S6,S7) and the positions of the boundaries between regions of different cell shapes. (B) Plot of the meridional intrinsic curvature $\kappa_s^0$ against $s$, defining additional fitting parameters $\kappa_1, \kappa_2, \kappa_3$ encoding the intrinsic curvatures of the different cell shapes and the additional positional fitting parameter $s_1 < s_2$. (C) Plot of the circumferential intrinsic curvature $\kappa_\phi^0$ against $s$, defining the additional fitting parameter $\kappa_4$ that defines yet another intrinsic curvature value for a cell shape. As in Ref. (S6), the parameter $\sigma$ regularising what would otherwise be discontinuities in $f_s^0, f_\phi^0, \kappa_s^0, \kappa_\phi^0$ is not fitted for.

**Fitting methods and details.** For fitting this model to the average shape of *Volvox* at a given developmental timepoint of inversion, similarly to Ref. (S6), we place fitting points $(R_i, Z_i)$ for $i = 1, 2, \ldots, N_{\text{fit}} = 100$ at equal spacing along the arclength of the meridian of this average shape. For a given set of parameter values, we solve the boundary value problem associated with Eqs. **7** of the main text using the `bvp4c` solver of MATLAB (The MathWorks, Inc.) or a custom implementation of arclength continuation based on the same solver. The solution of this boundary value problem defines a cross-section $(r(s), z(s))$, where $\mathrm{d}z/\mathrm{d}s = f_s \sin\psi$ (*Materials and Methods* of the main text). This yields points $(r_i, z_i)$ equally spaced along this cross-section, for $i = 1, 2, \ldots, N_{\text{fit}} = 100$. The fitting minimises the fit energy

$$E_{\text{fit}} = \left\{ \sum_{i=1}^{N_{\text{fit}}} \left[ (r_i - R_i)^2 + (z_i - Z_i)^2 \right] \right\}^{1/2} \qquad \textbf{[S60]}$$

over the fitting parameters. We perform this minimisation using a modification of the `fminsearch` function of MATLAB that uses the modified parameters for its underlying Nelder–Mead algorithm suggested by Ref. (S8) and that restricts the initial simplex used by the algorithm. The second modification avoids parameter values that are "too far way" from the initial guess and that might lead to the algorithm failing due to bifurcations that lie between the initial guess and these parameter values in parameter space. Further, we stopped the minimisation once $E_{\text{fit}}$ dropped below a certain value (Fig. 3N of the main text); empirically, we found that this avoids excessive variations of fit parameters. We fit the average shapes for the developmental timepoints $t_1 < t_2 < \cdots$ one after the other, using the fitted parameter values from the previous timepoints to obtain initial guesses.

***Fitting details for the different fits in the main text.*** For the fits discussed in the main text, the number of fitting parameters was reduced as follows: For the fits of constant $\kappa_1 = \kappa^0_{\text{post}} \in \{0, R^{-1}, r^{-1}_{\text{post}}\}$, the parameters that were fitted for at each timepoint were $f_2, f_3, f_4, f_5, \kappa_2, s_1, s_2, s_3, s_4$. Additionally, $f_1 = r_{\text{post}}/R, \kappa_3, \kappa_4$ were fitted only at the first timepoint and held constant for later timepoints. This assumption reduces the number of fitting parameters. For the final "free" fit, $f_1 = r_{\text{post}}/R, \kappa_1 = \kappa^0_{\text{post}}, \kappa_3, \kappa_4$ were additionally fitted for at each timepoint.

**Further discussion of the fitting results.** In this final subsection, we discuss the fitting results of the main text further. In particular, we must discuss the fact that we did not fit the parameters $f_1, \kappa_3, \kappa_4$ at each timepoint. Based on the observed cell shape changes (S7), no variation of $f_1$ is expected while the cells in the posterior are spindle-shaped. This is borne out by the "free" fit, which suggests that $f_1$ remains (approximately) constant until around the time of phialopore opening, when the cells in the uninverted posterior become pencil-shaped (S7), as discussed in the main text. Not fitting the parameters $\kappa_3, \kappa_4$ is *a priori* a simplifying assumption, because the preferred curvatures of the teardop-shaped cells and disc-shaped cell shapes that are observed in the anterior hemisphere during early and late inversion (S7) could be different. Could the worse fit energies for the fits with $\kappa^0_{\text{post}} \in \{0, R^{-1}\}$ (Fig. 3N of the main text) result from the different values of $r_{\text{post}}$ (inset of Fig. 3O in the main text) or $\kappa_3, \kappa_4$ fitted at the first timepoint? To exclude this possibility, we have run additional fits with $\kappa^0_{\text{post}} = 0$ using the values of $f_1 = r_{\text{post}}$ and/or $\kappa_3, \kappa_4$ used for $\kappa^0_{\text{post}} = r^{-1}_{\text{post}}$. These fits did not lead to better fit energies (Fig. S2). The fact that the variations of the fitted values of $\kappa^0_{\text{post}}$ and $r_{\text{post}}$ in the final "free" fit (Fig. 3O of the main text) are comparable to the differences between these values in the earlier fits is also consistent with this. All of this shows that these simplifying assumptions do not cause the worse fits, and hence that these result indeed from the "wrong" values of $\kappa^0_{\text{post}}$.

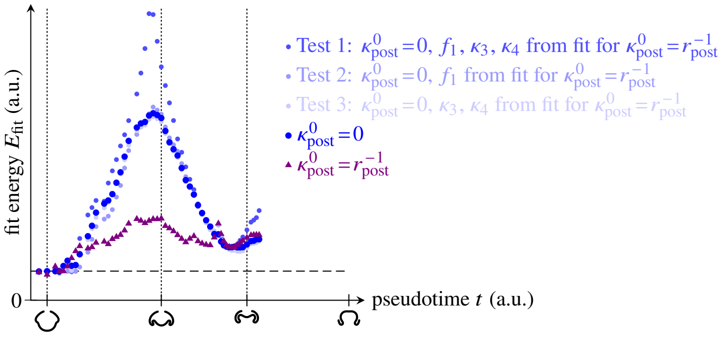


**Fig. S2.** Test of the fit results in the main text. Plot of the fit energy $E_{\text{fit}}$ against developmental pseudotime $t$: Repeating the fits for $\kappa^0_{\text{post}} = 0$ using the values of the parameters $f_1 = r_{\text{post}}$ and/or $\kappa_3, \kappa_4$ from the (better) fit with $\kappa^0_{\text{post}} = r^{-1}_{\text{post}}$ does not lead to better fits. The energies of the fits for $\kappa^0_{\text{post}} \in \{0, r^{-1}_{\text{post}}\}$ are repeated from Fig. 3N of the main text for comparison. Pseudotimepoints used for the discussion in Fig. 3D–L of the main text are highlighted by the shape insets on the horizontal axis.
